# Extracellular hyaluronate pressure shaped by cellular tethers drives tissue morphogenesis

**DOI:** 10.1101/2020.09.28.316042

**Authors:** Akankshi Munjal, Edouard Hannezo, Timothy J. Mitchison, Sean G. Megason

## Abstract

How tissues acquire complex shapes is a fundamental question in biology and regenerative medicine. Zebrafish semicircular canals form from invaginations in the otic epithelium (buds) that extend and fuse to form the hubs of each canal. We find that conventional actomyosin-driven behaviors are not required. Instead, local secretion of hyaluronan, made by the enzymes *ugdh* and *has3*, drives canal morphogenesis. Charged hyaluronate polymers osmotically swell with water and generate isotropic extracellular pressure to deform the overlying epithelium into buds. The mechanical anisotropy needed to shape buds into tubes is conferred by a polarized distribution of cellular protrusions, linked between cells, that we term cytocinches. Most work on tissue morphogenesis ascribes actomyosin contractility as the driving force, while the extracellular matrix shapes tissues through differential stiffness. Our work inverts this expectation. Hyaluronate-pressure shaped by anisotropic tissue stiffness may be a widespread mechanism for powering morphological change in organogenesis and tissue engineering.

## Introduction

Most organs including kidney, vasculature, lung, gut and heart begin as simple epithelia that then undergo morphogenesis to create elaborate shapes necessary for their function (Lubarsky and Krasnow, 2003). How this occurs is a fundamental question in biology with important implications in disease and regenerative medicine. Most research across tissues and species impute actomyosin networks as the central producers of mechanical forces that power cell deformations during tissue morphogenesis (Heisenberg and Bellaiche, 2013; Munjal and Lecuit, 2014). Actomyosin networks generate active contractile tension through the pulling of Actin filaments (F-Actin) by non-muscle Myosin II motor proteins (Myosin II) using ATP hydrolysis (Hartman and Spudich, 2012). The F-actin cortex is anchored to the plasma membrane and transmits forces across the tissue through cell adhesion (e.g. Cadherins and Integrins) (Lecuit et al., 2011). In this paradigm, developmental signals and biochemical information pattern tissue dynamics, such as bending, expansion and elongation, by locally activating actomyosin contractility and/or modulating cell adhesion, to drive cell behaviors, like apical constriction, division and rearrangement. Classic examples include apical constriction during the invagination of the presumptive mesoderm (Dawes-Hoang et al., 2005), and the planar polarization of cell-cell junction strength to control cell arrangements during convergence-extension (Bertet et al., 2004).

The extracellular matrix (ECM), that is attached to most tissues, has long been recognized as an important determinant of tissue mechanics by integrating tissue forces and by providing stiffness. The ECM gets its stiffness from fibrous protein polymer networks such as fibronectins, laminins and collagens. In this regime, the key mechanical aspect of the ECM is its elasticity and ability to transmit tensile force across cells in an epithelium. During tissue morphogenesis, the ECM can create anisotropic shapes by resisting isotropic forces through differential stiffness (Dzamba and DeSimone, 2018). For instance, the circumferentially aligned collagen fibers in the ECM of the *Drosophila* egg constrain isotropic tissue growth to facilitate elongation in the anterior-posterior axis (Crest et al., 2017). The ability of collagen fibers to assemble into gels with high tensile strength, especially after cross-linking, is key to the regime where the ECM plays a passive, elastic role in morphogenesis (Chaudhuri et al., 2020).

Hyaluronic acid (HA) is another widespread ECM component, but its physical properties differ greatly from collagens, fibronectin and laminins. HA is a long, flexible polysaccharide with a high density of carboxylate groups that are mostly negatively charged and balanced by sodium ions under physiological conditions. As a result of this chemistry, HA polymers tend to swell with water and form viscoelastic hydrogels with a low mass fraction, low tensile strength but high propensity to generate compressive osmotic forces by swelling (Cowman et al., 2015; Toole, 1981). Despite its abundance, the role of HA in tissue morphogenesis due to its biophysical properties has not been fully investigated. Rather, the biophysical HA literature is dominated by medical applications in restorative surgery, drug delivery and other areas (Borzacchiello et al., 2015).

We investigated how morphogenic forces are generated to deform a simple epithelium in an understudied yet exemplary system common to all vertebrate organisms, the semicircular canal (SCC) development (Higuchi et al., 2019). All jawed vertebrates have three mutually orthogonal SCCs in each inner ear whose well-conserved shape is required for their function of sensing balance and acceleration (Groves and Fekete, 2012) (Figure 1B). Head movement results in fluid movement in the canals, which is transduced by the hair cells to sense motion. Morphogenesis of the SCCs is among the most geometrically complex and precise events in vertebrate development (Figure 1A). The vertebrate inner ear forms from a thickening of the embryonic ectoderm located lateral to the hindbrain, called the otic placode (Whitfield, 2015). The otic placode cavitates or invaginates to form a tight epithelial fluid-filled structure called the otic vesicle (OV) (Whitfield, 2015). The single-layered OV then undergoes a dramatic topological change to form the SCCs (Alsina and Whitfield, 2017) (Figure 1A). The mechanisms underlying SCC morphogenesis remain poorly understood due to their inaccessibility in most model organisms. Zebrafish lack middle and outer ears, and the inner ear forms before ossification of the skull, which encases the inner ear, making it both optically and physically accessible, and thereby an excellent model organism to study SCC morphogenesis.

**Figure 1.**
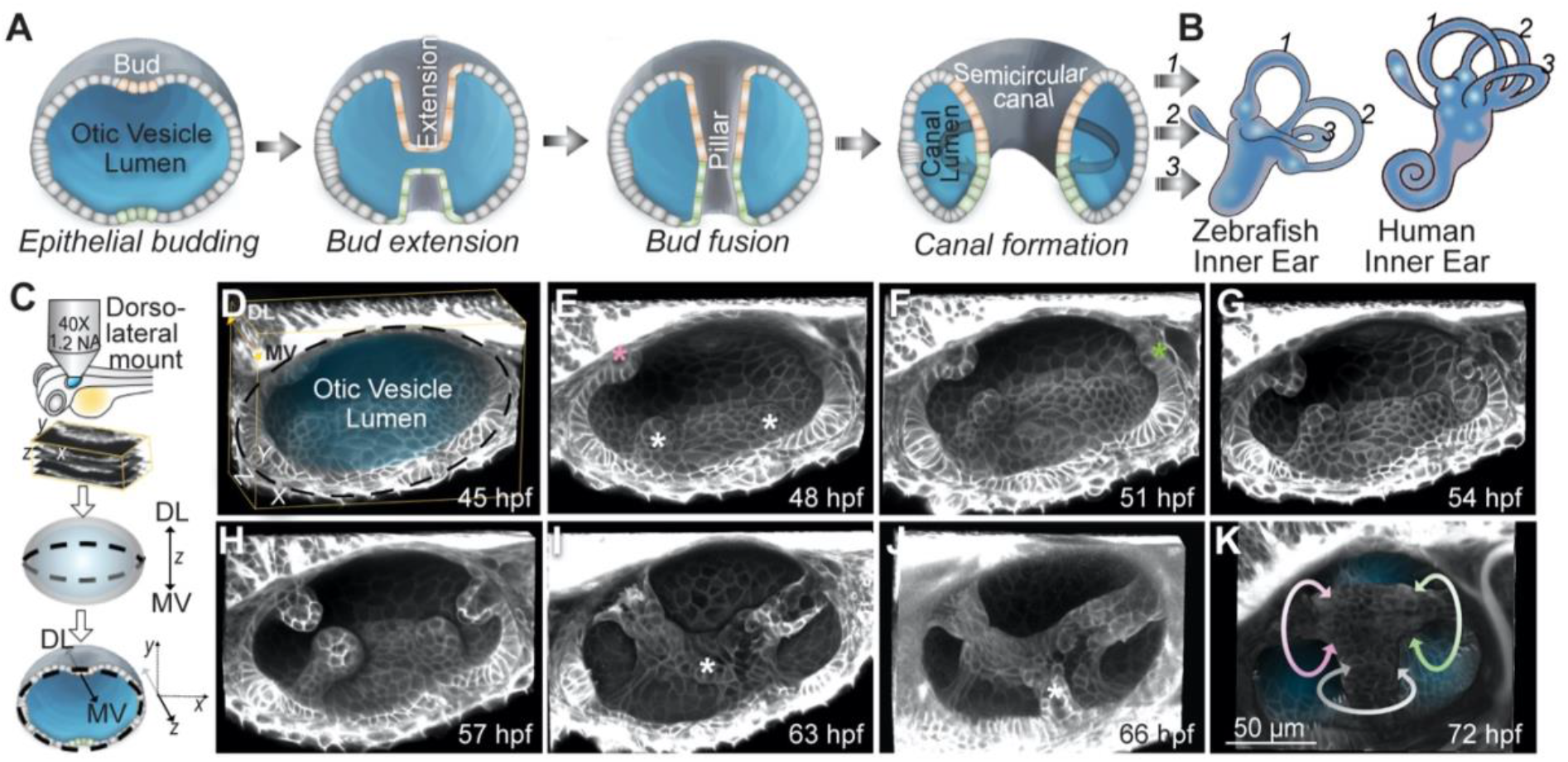
*In toto* imaging of the developing zebrafish inner ear reveals multi-scale dynamics during semicircular canal morphogenesis. (**A**) Illustrations showing the formation of a SCC. (**B**) Illustrations of the inner ears from adult zebrafish and human comparing the conserved structures of the SCCs. Anterior, posterior and lateral SCCs are labelled as 1, 2 and 3 respectively. (**C**) Workflow for zebrafish larva mounting, image acquisition and visualization. (**D-K**) 3D rendered otic vesicles (OVs) at select time points using *Tg(βActin:membrane-Citrine)*. Anterior to the left and dorso-lateral (DL) into plane of view. Lateral, anterior and posterior buds are marked by white, pink and green asterisks respectively. The buds extend (G and H), and fuse (I). Ventro-lateral and ventral buds form and extend (marked by white asterisks) (I and J). Bud fusion demarcates the hubs for the anterior, posterior and lateral SCC (marked by pink, green and white circular arrows respectively) (K). Scale bar, 50 μm.

Using *in toto* imaging of transparent zebrafish embryos, perturbation approaches and biophysical modelling, we discovered unexpected roles of the ECM and actomyosin networks in SCC morphogenesis. Patterned cells in the otic epithelium locally generate extracellular HA, which in turn actively reshapes the tissue through osmotic swelling. Thus, the driving force is ultimately ECM-derived osmotic pressure rather than actomyosin contractility. Actomyosin networks, instead, play a role in directing anisotropic morphogenesis through differential tissue stiffness provided by the intercellular tethers we term cytocinches.

## Results

### Long-term imaging of the developing zebrafish inner ear reveals multi-scale dynamics during semicircular canal morphogenesis

All prior studies on semicircular canal development have been done with either low resolution live microscopy or static imaging by fluorescence, paint-fills or micro-CT images (Chang et al., 2004; Groves and Fekete, 2012; Nishitani et al., 2017; Whitfield et al., 1996). To perform long-term in toto imaging of zebrafish SCC development, we take advantage of transgenic zebrafish expressing a bright membrane-localized fluorescent protein, thereby minimizing photo-toxicity and photo-bleaching (Swinburne et al., 2018). Instead of anesthetizing with tricaine, which slows inner ear development with long-term exposure, we immobilized larvae with alpha-Bungarotoxin (aBt) that allows normal development (Swinburne et al., 2015). Larvae were mounted dorso-laterally as shown in Figure 1C, in a cast agarose mount that secures the head and the yolk, and positions the ear just beneath the cover slip. This set up allows long-term imaging of the SCC development at sub-cellular and high temporal resolution.

SCCs form from the topological remodeling of the OV sandwiched between the skin and the hindbrain and surrounded by mesenchymal cells (Figure 1D, Figure S1A and Movie S1). Morphogenesis initiates when cells from six different regions of the otic epithelium sequentially form buds projecting into the lumen (Figures 1E, 1F, 1I, 1J, S1A, S2C and Movie S1). Cells in lateral (antero-lateral and postero-lateral) and anterior regions form buds first (45 hpf), followed by cells in the posterior region (51 hpf), and lastly in the ventro-lateral and the ventral regions (63 hpf) (Figures 1E, 1F, 1I, 1J, S1A-S1C, and Movie S1). The six buds extend anisotropically in their longitudinal axis (Figures 1F-1J, S1B-S1F, and Movie S1). Topology change occurs when the buds fuse to one another to form three pillars through the OV lumen demarcating the hubs of the future anterior, posterior and lateral SCCs (Figures 1I-1K and Movie S1).

### Stereotypical morphogenic behaviors are not responsible for SCC morphogenesis

To investigate the mechanisms underlying SCC development, we tested existing models for morphogenesis, including elevated proliferaton of bud cells compared to the neighboring cells (Figure 2K). Localized increase of cell proliferation has been observed in nascent epithelial buds during branching morphogenesis in a number of organs including kidney, salivary gland and lung (Varner and Nelson, 2014). Using a proliferative marker, 5-ethynyl-2′-deoxyuridine (EdU) staining, we find a low rate of proliferation during SCC morphogenesis (45 – 54 hpf) with 25% of cells in the lateral region of the OV, and less than 10% of the bud forming cells in S phase (Figures 2A and 2C). Inhibiting S phase with Hydroxyurea and Aphidicolin between 48-54 hpf reduced EdU incorporation down to 10% in the lateral OV and to 0% in bud cells, yet their morphogenesis continued normally with a slight delay (Figures 2B-2D). We conclude that localized cell proliferation is not responsible budding.

**Figure 2.**
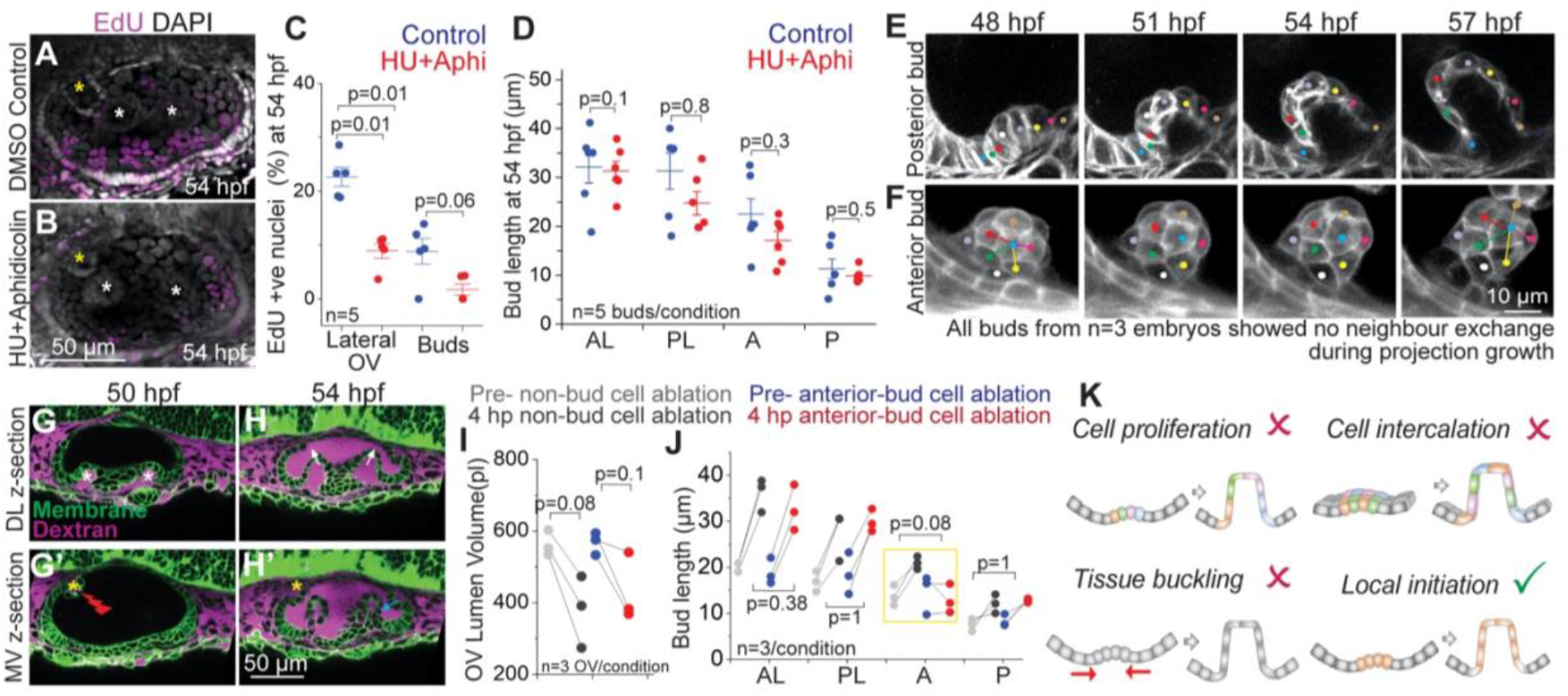
Stereotypical morphogenic behaviors are not responsible for SCC morphogenesis. **(A and B**) 3D rendered representative examples of OVs with DAPI and EdU staining marking all nuclei (grey) and nuclei in S phase (purple) respectively, in control (A), and Hydroxyurea (HU)+Aphidicolin treated larvae (B) at 54 hpf. Lateral buds are marked by white asterisks and anterior bud is marked by yellow asterisks. Anterior to the left and DL into the plane of view. Scale bar, 50 μm. (**C**) Individual data points and mean±s.d. of percentage of EdU positive nuclei in the lateral region of the OV, and only in the buds. ‘n’ denotes the number of OVs. p values as labelled (Mann Whitney- U test). (**D**) Individual data points and mean±s.d. of bud lengths in control and HU+Aphidicolin treated embryos at 54 hpf. In the absence of buds, lengths correspond to cell lengths. ‘n’ denotes the number of buds. p values as labelled (Mann Whitney- U test). (**E and F**) 2D sections (E) and 3D rendering (F) of a posterior and an anterior bud respectively from membrane-labeled transgenic larva at select time points with individual cells tracked (colored dots). Scale bar, 10 μm. (**G and H**) 2D sections from two different z-depths of an ablated OV (marked by red flash) using *Tg(βActin:membrane-Citrine)* (green) and Texas-red dextran (magenta) in the periotic space. Lateral buds (marked by white asterisks) can be seen in the DL section (L). Anterior bud (marked by yellow asterisky) can be seen in the MV section (L’). Dextran dye enters the OV lumen upon ablation (M-M’). Lateral buds continue to extend in the ablated OV (marked by white arrows) (M’). Posterior bud also forms and extends (cyan arrow). Anterior bud ablation blocks its extension (yellow asterisk) (M’). Scale bar, 50 μm. (**I**), OV lumen volume in control and experiment before and after ablation. ‘n’ denotes the number of OVs per condition. p values as labelled (Mann Whitney- U test). (**J**) Bud lengths in control and experiment larvae before and 4 hours post (hp) ablation. ‘n’ denote number buds measured per condition. p values as labelled (Mann Whitney-U test). (**K**) Illustrations show the models tested for budding morphogenesis.

We next investigated whether cell-rearrangement based convergence-extension drives otic epithelial bud extension (Figure 2K). Convergence-extension is a widely employed mechanism for tissue elongation during embryonic development (Wallingford et al., 2002). During *Drosophila* salivary gland morphogenesis, cells exhibit circumferential convergence and radial extension to form a narrow tube from a round and flat epithelium (Sanchez-Corrales et al., 2018). Given the geometrical similarity, we tested this model in the otic epithelium by tracking cells in the morphogenic regions. We observe that as buds extend, cells adjacent to the bud become part of the bud (Figure 2E). However, both the cells adjacent to and within the bud, maintain their neighbors during bud extension, thereby refuting a cell-rearrangement based model for budding morphogenesis (Figures 2E and 2F).

We next examined tissue-scale mechanisms such as buckling in SCC morphogenesis (Figure 2K). Compressive stresses from the differential growth of apposed tissues can cause tissues to buckle or fold (Nelson, 2016). Buckling drives villi formation in the small intestine (Shyer et al., 2013) and branching in the airways of the lung (Nelson, 2016). Besides tissue growth, osmotic pressure from interstitial and lumenal fluids can also apply stresses on the surrounding epithelium (Chan et al., 2019; Navis and Bagnat, 2015), including in the OV (Mosaliganti et al., 2019). As reported above, cell proliferation is low during SCC morphogenesis and blocking it does not affect bud extension (Figures 2A-2D). To reduce lumen pressure, we used a laser-mediated targeted ablation of single cells in the OV to disrupt the epithelial barrier causing loss of lumen volume and pressure (Mosaliganti et al., 2019) (Figure 1G and 1I). Ablation-mediated loss of lumen pressure neither affected initiation of new buds nor the extension of existing buds (Figures 1H and 1J). Together these data are inconsistent with a buckling-based model for morphogenesis (Figure 2K). Interestingly, ablation of one or two bud cells was sufficient to block extension of that bud, while the adjacent non-ablated buds continued morphogenesis normally (Figures 2G-2J), showing that morphogenesis of each bud is locally initiated.

### Semicircular canal morphogenesis requires patterned expression of hyaluronan synthesis enzymes *ugdh* and *has3*

We next examined a role for the extracellular matrix (ECM) in the local initiation of each bud (Figure 2K). In mouse gut explants, mesenchymal condensates have been shown to align the surrounding Collagen I fibers causing the overlying epithelium to bend (Hughes et al., 2018). In the frog inner ear, hyaluronic acid (HA) was shown to be required for SCC formation and suggested to act as a propellant to drive bud extension (Haddon and Lewis, 1991). HA is a secreted, unbranched and non-sulfated glycosaminoglycan composed of repeating disaccharide subunits. HA is synthesized by the enzymes uridine 5′-diphosphate dehydrogenase (*ugdh)*, which participates in subunit synthesis, and hyaluronan synthase (*has)* that polymerizes subunits at the cell membrane to extrude out chains directly into the extracellular space (Vigetti et al., 2014). Which cells are responsible for HA-production and how HA generates morphological change during SCC formation remains unclear.

We investigated whether HA synthesis enzymes are expressed by the mesenchymal cells that surround the OV, or by the cells of the otic epithelium, using multiplexed fluorescent *in situ* hybridization (Choi et al., 2018). We observed local and uniform expression of *ugdh* and *has3* in the bud cells of the OV (Figures 3A-3C, S2A and S2B). In contrast, Collagen 2 (*col2a1a*), a major component of the ECM, is not exclusive to the buds and instead expressed in all the cells of the OV (Figures 3A-3C and S2C), while *has2*, known for its role in cardiac morphogenesis (Patra et al., 2011), is not expressed in the OV (C. Thisse, 2005). Moreover, in contrast to *col2a1a, ugdh* and *has3* expression were not observed after the buds fused (Figures S2A-S2C). We conclude that *ugdh* and *has3* are locally, uniformly and transiently expressed in the bud cells of the OV during morphogenesis.

**Figure 3.**
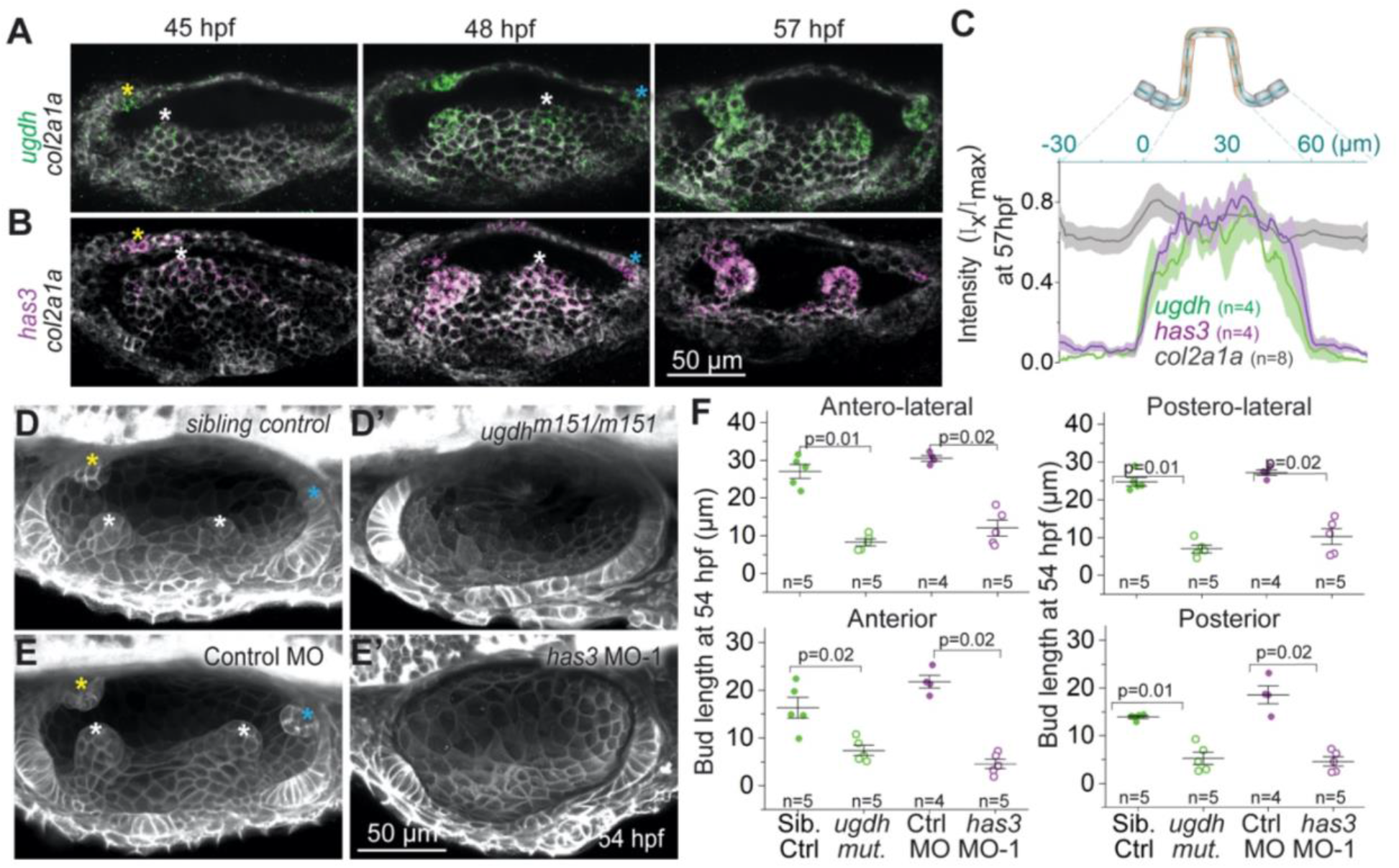
Semicircular canal morphogenesis requires patterned expression of hyaluronan synthesis enzymes *ugdh* and *has3*. (**A and B**) Maximum intensity projections of OVs at select time points stained with multiplex *in situ* probes against *ugdh* (green) and *col2a1a* (white) (A), and *has3* (purple) and *col2a1a* (white) (B). The z-volume is different across time points to capture all the buds, and hence the contrast of each time point is individually set for better visualization. Scale bar, 50 μm. (**C**) Mean intensities±standard error (s.e.) of various genes across the illustrated region of interest (ROI) in the anterior bud at 57 hpf. (**D and E**) Representative examples of 3D-rendered OVs at 54 hpf from sibling control and *ugdh*^*m151/m151*^ mutant larvae labelled with *membrane-NeonGreen* mRNA (D), and *Tg(βActin:membrane-Citrine)* embryos injected with control morpholino (MO) and *has3*-specific MO-1 (E). Buds in controls are marked by asterisk (white, yellow and cyan for lateral, anterior and posterior buds respectively). Genetically perturbed larvae have no buds. Anterior to the left and DL into the plane of view. Scale bar, 50 μm. (**F**) Individual data points and mean±standard deviation (s.d.) of bud lengths in controls and genetic perturbations at 54 hpf. In the absence of buds, bud lengths correspond to cell lengths. ‘n’ denotes the number of buds measured per condition. p values as labelled (Mann Whitney- U test).

We next tested the function of these enzymes in SCC morphogenesis. A point mutation that disrupts *ugdh* (Driever et al., 1996; Walsh and Stainier, 2001) blocked OV budding at 54 hpf (Figures 3D and 3F) and caused aberrant bud morphologies at 72 hpf (Figure S2D). Morpholino-mediated knockdown of *has3* (Ouyang et al., 2017) also abrogated SCC formation with no budding at 54 hpf (Figures 3E and 3F) and aberrant bud morphologies at 72 hpf (Figures S2E and S2F). We conclude that *ugdh* and *has3* are required for zebrafish SCC formation. The requirement of HA for SCC development is potentially conserved across species as specific expression of *has2* is observed in the SCCs during mouse development (Tien and Spicer, 2005).

### The ECM of the buds is rich in hyaluronan and dense

We investigated the mechanical basis of HA-dependent morphogenesis. In physiological conditions, HA is made as long charged polymers (0.5-6MDa) behaving as a polyelectrolyte, which imbibes and retains large amounts of water to a form a viscoelastic solution (Cowman et al., 2015). Owing to its unique biophysical properties, HA is directly involved in tissue homeostasis and biomechanics in adult organisms (Dicker et al., 2014). However, applications of these properties in tissue morphogenesis during development are not well-understood.

We checked the localization of endogenous HA using a HA- binding protein (HABP) (Kohda et al., 1996). During bud formation, HABP is restricted to and fills up the extracellular space beneath the buds (hereafter referred to as the bud-ECM) (Figures 4A-4C, 4E and S3A). In contrast, Collagen2 was detected in the ECM surrounding the entire OV with slightly higher expression in the bud-ECM (Figures 4A-4C, 4E and S3A). We perturbed HA by injecting hyaluronidase, an enzyme that breaks down HA polymers into oligomers, in the space surrounding the otic vesicle (periotic space). Hyaluronidase treatment completely inhibited HA accumulation in the bud-ECM (Figures 2D and S3B) showing that local deposition of HA depends on its polymeric properties.

**Figure 4.**
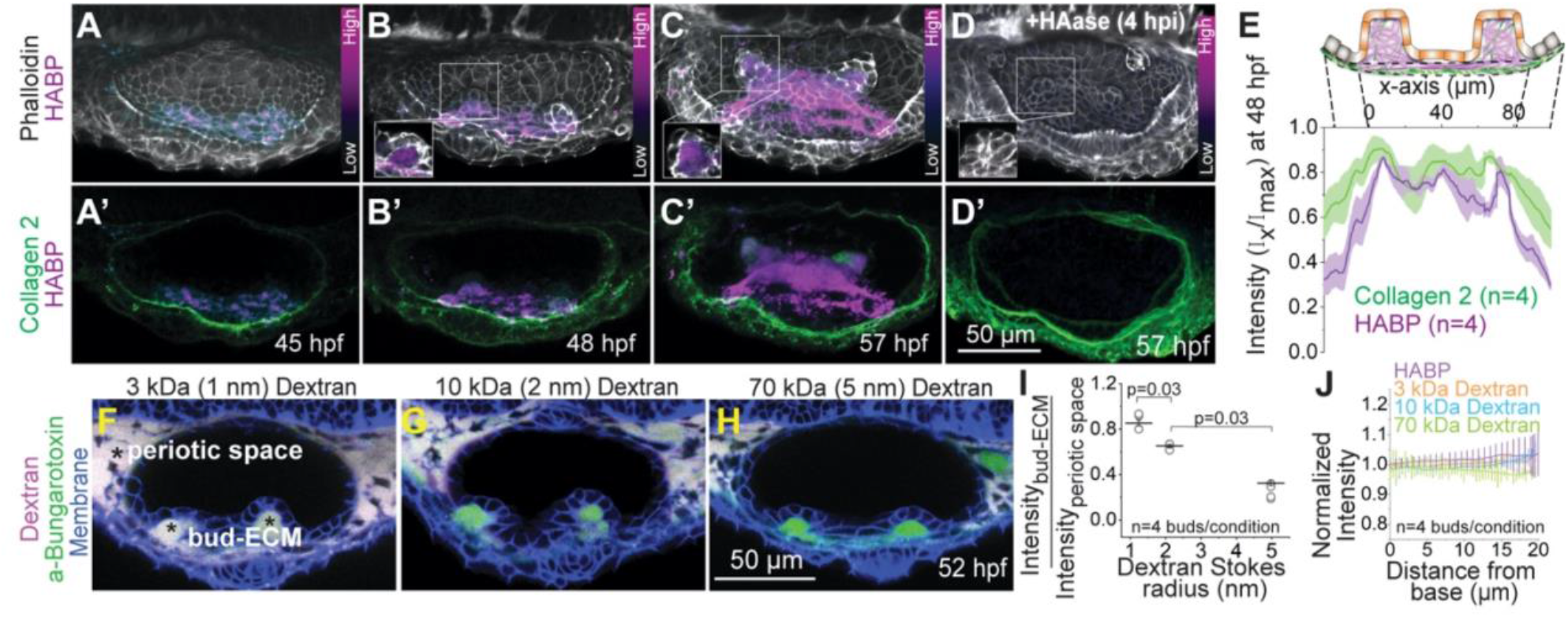
The ECM of the buds is rich in hyaluronan and dense. (**A-D**) 3D rendered OVs showing HA, F-actin and Collagen 2 staining using HA-Binding Protein (HABP), Phalloidin and Anti-Col2a1a respectively, at select time points in uninjected (A-C) and hyaluronidase treated larvae (D). Anterior to the left and DL into the plane of view. The contrast of each time point is individually set to capture the dynamic range of HABP. Insets show 2D sections of the antero-lateral buds. Scale bar, 50 μm. ‘n’ denotes the number of buds. (**E**) Mean intensities±s.e. of various stains across the illustrated ROI in the lateral buds at 48 hpf. (**F-H**) 2D sections showing percolation of dextran from the periotic space into the bud-ECM 2 hours post injection (hpi). Different sizes of Texas-red dextran (in magenta)-3 kDa (F), 10 kDa and 70 kDa (H) with approximate Stokes radii 1 nm, 2 nm and 5 nm respectively, were co-injected with aBt (green) in *Tg(βActin:membrane-citrine)* larvae (blue). aBt colocalizes with all three sizes of dextran in the periotic space (white). Contrast is same across larvae. Scale bar, 50 μm. (**I**) Individual data points and mean±s.e. of fluorescent intensities of different sizes of dextran in the bud ECM normalized to their intensities in the periotic space. ‘n’ denotes the number of buds per condition. p values as labelled (Mann Whitney- U test). (**H**) Mean±s.e. of the normalized fluorescent intensities of different sizes of dextran and HABP in the bud ECM from the base to the tip (as show in the illustration Figure S3C). ‘n’ denotes the number of buds per condition.

To probe the physical properties of HA, we measured the percolation of different sizes of fluorescent dextran (molecular weights 3 kDa, 10 kDa, and 70 kDa and approximate Stokes radii 1.3, 2.4 and 5 nm respectively (Granath, 1958)) into the bud-ECM when injected in the space periotic space (Figures S3C and 4F-4H). Fluorescent labelled aBt was co-injected with dextran as a control. After 2 hours, aBt equilibrated in the bud-ECM relative to the periotic space, while 85% of 3 kDa, 65% of 10 kDa, and only 30% of 70 kDa dextran percolated in the bud-ECM (Figures 4F-4I). Low percolation of large dextrans shows that the HA-rich bud-ECM is dense with a pore size in the nanometer range. We find the intensity of HABP and percolation of dyes to be uniform within the bud-ECM suggesting uniform distribution and structure of the HA-rich ECM (Figures S3C and 4J).

### Hyaluronate drives cellular and tissue morphogenesis through isotropic pressure

We investigated the effect of the HA-rich bud-ECM on the overlying otic epithelial cells. HA can apply pressure on the surrounding tissues due to osmotic-swelling (Toole, 1981). As the buds form and extend, the volume of the bud-ECM increases (Figures 5A, 5C and Movie S2). Inhibition of HA with hyaluronidase treatment led to a significant reduction of bud-ECM volume (Figures 5B and 5D), and in agreement with previous results, arrested bud extension (Figures 5B and 5E) (Geng et al., 2013). We tested if the swelling of the HA confined in the bud-ECM applies hydrostatic pressure on the overlying bud cells in the form of cell strain (Figure 5F). Consistent with our hypothesis, during bud extension, the HA- producing bud cells thin in the radial axis and stretch in the circumferential and longitudinal axes, while maintaining their volume (Figures 5C, 5G, 5H and Movie S3). Strikingly, hyaluronidase treatment causes loss of cell stretching in the buds (Figures 5G and 5I). Together these data show that hyaluronate-pressure remodels the HA-producing cells to power cell stretching and epithelial budding (Figure 5F).

**Figure 5.**
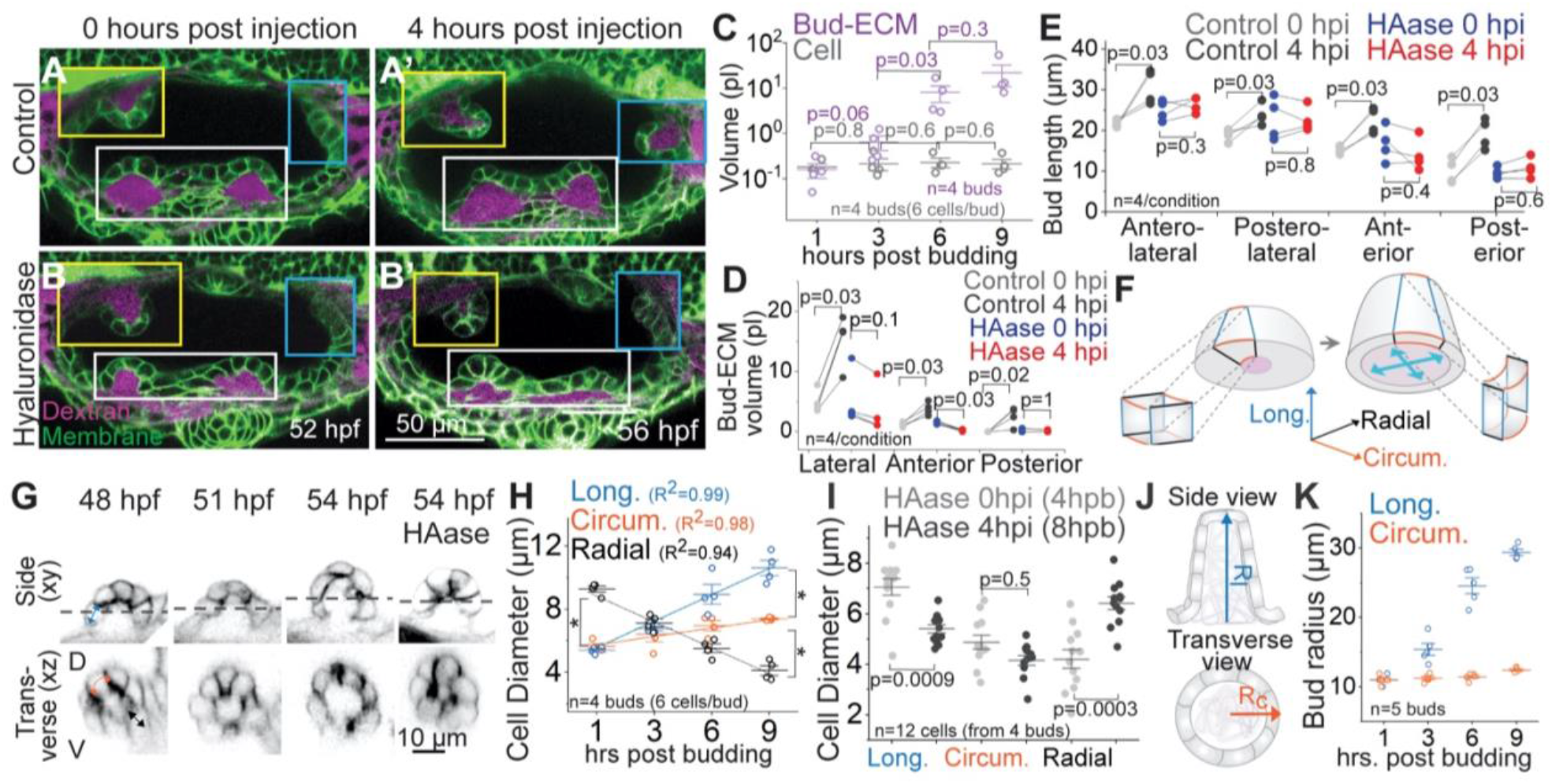
Hyaluronate drives tissue and cellular morphogenesis through isotropic pressure. (**A and B**) 2D sections of representative examples from controls (A) and HAase injected (B) larvae at 0 and 4 hours post injection (hpi) using *Tg(βActin:membrane-Citrine)* (green) and Texas-red dextran (magenta) in the periotic space. Lateral, anterior and posterior buds from different z-depths are framed in white, yellow and cyan respectively. (**C**) Individual data points and mean±s.d. of bud-ECM and cell volume across time. Each data point for cell volume is an average from 6 cells in a bud. ‘n’ denotes the number of buds per condition. p values as labelled (Mann Whitney- U test). (**D and E**) Bud-ECM volume (D) and bud lengths (E) in control and HAase injected larvae at 0 and 4 hpi. In the absence of buds, lengths correspond to cell lengths. ‘n’ denotes the number of bud-ECMs or buds respectively. p values as labelled (Mann Whitney- U test). (**F**) Illustration showing the longitudinal (blue), circumferential (organa) and radial (black) axes of cells during budding. Deposition of HA in the bud-ECM is shown in purple. Cyan arrows show hydrostatic pressure from HA-swelling. Notice the change in aspect ratio of the cells before and after HA-swelling. (**G**) Side and transverse sections of an anterior bud from an uninjected larva at select time points, and HAase treated larva at 54 hpf, using *Tg(βActin:membrane-Citrine)*. Dotted line in side view (xy) marks the *y* position for transverse view (xz). (**H**) Individual data points and mean±s.d. of cell diameters across time (hours post budding, hpb). *denotes p=0.03 (Mann Whitney- U test). Each data point is an average from 6 cells per bud. ‘n’ denotes the number of buds. (**I**) Individual data points and mean±s.d. of cell diameters in hyaluronidase-treated larvae at 0 and 4 hpi. Each data point is the diameter of a single cell. ‘n’ denotes the number of cells. p values as labelled (Mann Whitney- U test). (**J**) Illustration showing the longitudinal (R_1_) and circumferential radius (R_C_) of the bud. (**K**) Individual data points and mean±s.d. of bud radii across time. ‘n’ denotes the number of buds.

We next investigated how buds can grow anisotropically in the longitudinal axis. Time-lapse observation showed that cell stretching is greater in the longitudinal than the circumferential axis (Figure 5H and Movie S3). Similarly, buds first have an isotropic aspect ratio (1:1, longitudinal: circumferential radius) between 1-3 hours post budding (hpb), and then extend only in the longitudinal axis to acquire an anisotropic aspect ratio (2.5:1) (Figure 5K and Movie S3). We considered if anisotropic bud extension is mediated by HA. HA is comprised of long flexible polymers in solution and drives morphogenesis through hydrostatic pressure, which by Pascal’s law is spatially uniform within a static fluid. Moreover, the HA synthesis genes, *has3* and *ugdh*, are uniformly expressed across the bud epithelium (Figure 3C), and the intensity of HABP and percolation of dyes is uniform within the bud-ECM suggesting isotropic structural properties of the matrix (Figure 4J). Taken together, these data suggest that HA alone cannot be responsible for anisotropic bud extension.

### Tension-rich cellular tethers resist hyaluronate-pressure

We examined whether the bud epithelium itself provides the anisotropic stiffness needed to shape the isotropic hyaluronate pressure. Epithelial cells can alter their stiffness by increasing actomyosin network assembly to generate high internal tension (Lecuit et al., 2011; Tee et al., 2009). Using live reporters, we find that during bud extension, both F-actin and myosin II accumulate on the lateral and basal (ECM-facing) cell surfaces relative to the apical surface (OV lumen-facing) (Figures S4A-S4D and Movie S4). To better resolve this accumulation, we used mosaic membrane labelling and observed dynamic actomyosin-rich protrusions beneath the basal surface of adjacent cells (Figures 6A, 6B, S4E, S4F and Movie S5). Protrusive activity is commonly observed in migratory epithelia and is known to provide traction forces through cell-ECM interactions (Mayor and Etienne-Manneville, 2016). Protrusions in some non-migratory epithelia have also been observed with implications in inter-cellular communication either chemically (e.g. morphogen signaling by cytonemes in a number of tissues (Ramírez-Weber and Kornberg, 1999)) or mechanically (e.g. filopodial protrusions that align and anchor converging tissues (Chauhan et al., 2009; Jacinto et al., 2000; Zhang et al., 2020)).

**Figure 6.**
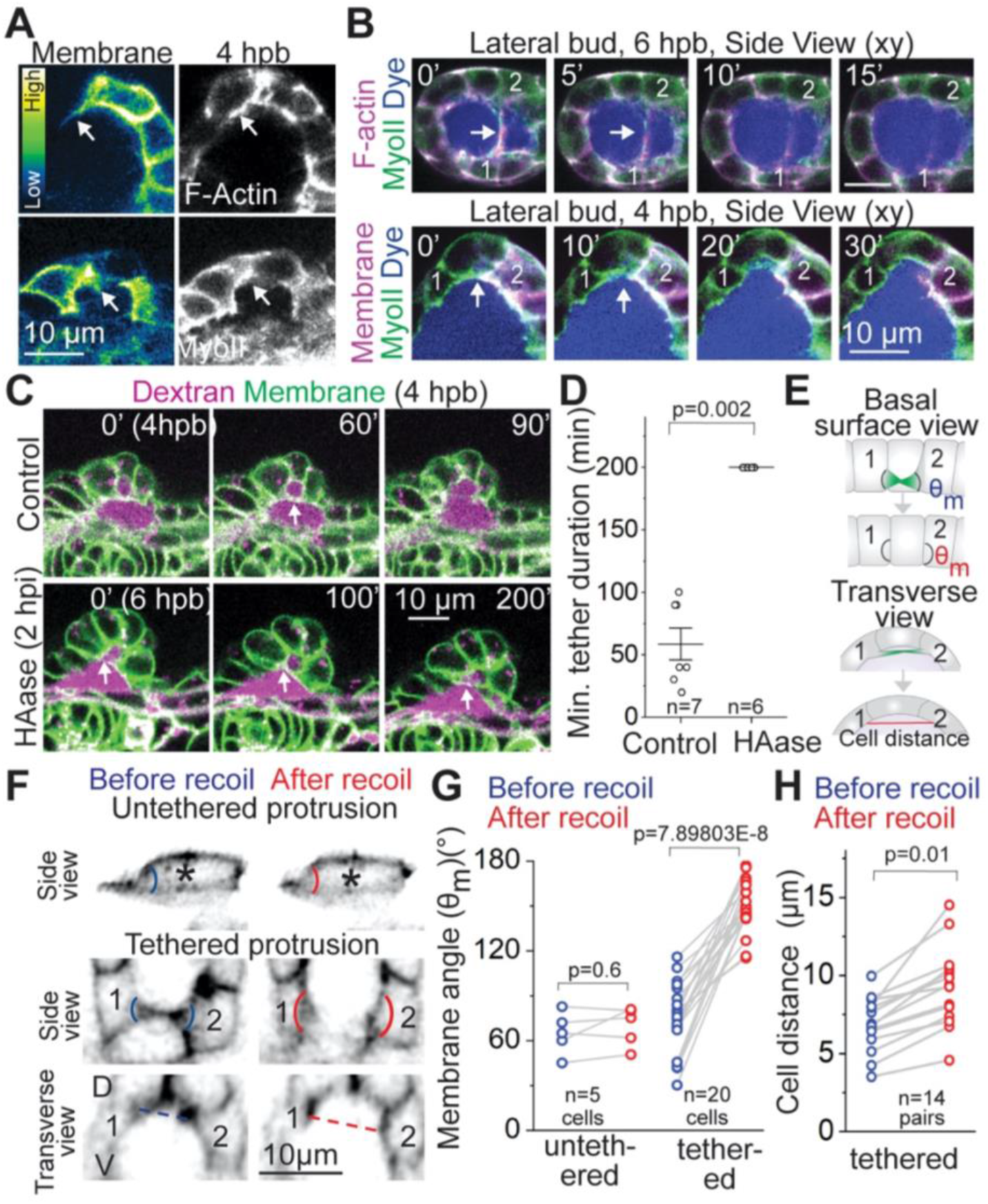
Tension-rich cellular tethers resist hyaluronate-pressure. (**A**) 2D sections of lateral buds showing protrusive activity with mosaic membrane labelling, and the corresponding localization of F-actin (top) and Myosin II (bottom). Scale bar, 10 μm. (**B**) 2D sections showing cells (1 and 2) with tethered protrusions (white arrow) in lateral buds using either F-actin (top) or mosaic membrane labelling (bottom) in magenta and Myosin II (green) reporters. Alexa Flour-647 conjugated aBt (blue) is injected in the periotic space. Scale bar, 10 μm. (**C**) 2D sections of lateral buds from controls (top) and HAase (bottom) injected larvae using *Tg(βActin:membrane-Citrine)* and Texas-red dextran in the periotic space. Tethered protrusions (marked by white arrow) retract in control (C) but not in HAase treated larva (D). Scale bar, 10 μm. (**D**) Individual data points and mean±s.d. of the minimum duration of tethered protrusions in control and HAase-treated larvae. n denotes the number of tethers. p values as labelled (Mann Whitney- U test). (**E**) Illustration showing a tethered protrusion (green). Adjacent membrane angles (θ_m_) are labelled in the basal surface view. Cell distance after tether recoils is marked by red line in transverse view. (**F**) Side and transverse views of 2D sections showing tethered and untethered protrusion retraction with membrane labelling. The cell forming an untethered protrusion (top) is marked with an asterisk. The cells forming tethered protrusions (bottom) are tracked with numbers 1 and 2. Adjacent membrane angles and adjacent cell distance are shown in blue and red before and after retraction respectively. Scale bar, 10 μm. (**G-H**) Adjacent membrane angles (G) and cell distance (H) before and after recoil. ‘n’ denotes the number of cells. p values as labelled (Mann Whitney- U test).

Given that protrusions occur beneath adjacent cells rather than through the bud-ECM (Movies S5 and S6), we tested if they provide inter-cellular mechanical communication. Strikingly, some protrusions from neighboring cells were tethered to each other and recoiled after spontaneous breaks (Figures 6B, 6C, 6F, S4F and Movie S6). Recoil was not observed in hyaluronidase treated embryos resulting in a longer tether duration and showing that HA swelling causes tethers to break (Figure 6C and 6D). Akin to laser ablation-induced recoil of cell-cell apical junctions to probe tension (Fierro-González et al., 2013; Rauzi and Lenne, 2015), HA pressure-induced recoil of tethered protrusions in control embryos served as a natural tension probe (Figure 6E and Movie S6). Recoil of tethers was associated with significant widening of adjacent membrane angles and an increase in neighboring cell distance (Figure 6F-6H and Movie S6). In contrast, untethered protrusion retraction did not associate with membrane angle change (Figure 6F and 6G). These measurements show that assembly of tension-rich tethered cellular protrusions provide resistance to ECM pressure. We therefore termed these protrusions “cytocinches”.

### Anisotropic distribution of cytocinches drives bud-to-tube transition

We tested whether the cytocinches are responsible for the anisotropy in bud shape. During bud extension, cytocinch tether numbers progressively increased and showed an orientation that was strongly biased towards the circumferential axis (Figures 7A, 7B and Movie S7). Strikingly, cytocinch anisotropy (number*orientation) correlated with the anisotropic aspect ratios of the bud (Figure 7C). If cytocinches resist turgor pressure from HA, this directional bias could explain the conversion of buds into tubes. To further explore this hypothesis, we built a theoretical model to examine if a cytocinch-based bias in resistance in the circumferential axis can result in bud extension in the orthogonal (longitudinal) axis (Figures 7D and S5). A 2D vertex model (Alt et al., 2017; Hannezo et al., 2014) representing a longitudinal “slice” of the epithelial monolayer enclosing a gradually increasing volume of ECM was used (See Methods for further details). The equilibrium configuration via energy minimization for a given volume was then calculated. Geometrical parameters and epithelial surface tension were estimated using membrane image data and actomyosin localization respectively (Figure S4A-S4D and Methods). The only free parameter was the inward tension from cytocinches applied to the basal vertices (Figures 7E, S5A-S5D and Methods). Consistent with our hypothesis, increasing the tension of circumferential cytocinches increased the anisotropy of the buds (Figures 7E, 7F, S5E, S5F and Movie S8). Interestingly, for intermediate values of cytocinch tensions, the bud short-axis remained roughly constant during 3-fold volume inflation, while the long-axis increased 3-fold (Figures 7F and S5E), as observed experimentally (Figure 5K). A good fit for bud anisotropy can be reached in simulations either by slowly increasing volume to a given level or simply starting at that level (Figures S5F and S5G), corroborating our conclusion that the bud-ECM is important for bud swelling, but not for its anisotropy. On the other hand, a progressive increase in cytocinch number was required for a good fit with experimental data, otherwise the model would overestimate early bud anisotropy (Figures S5F and S5H), corroborating the observed scaling between cytocinch and bud anisotropies (Figure 7C). Furthermore, the model predicted a relaxation of the bud to a hemispherical shape upon removal of cytocinch tension (Figure 7G and Movie S8).

**Figure 7.**
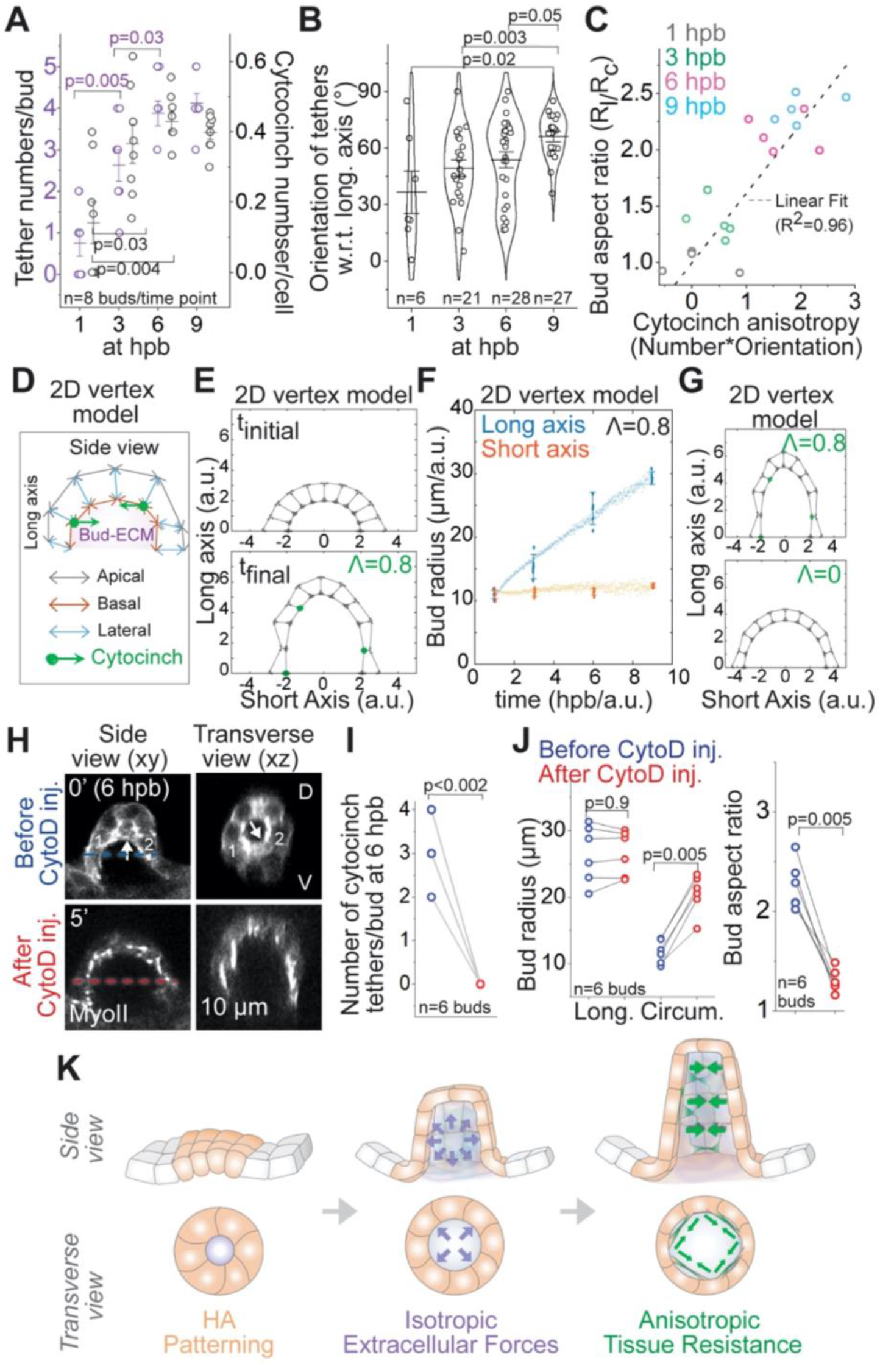
Anisotropic distribution of cytocinches drives bud-to-tube transition. (**A**) Individual data points and mean±s.d. of the number of tethers per bud (left axis, purple) and the number of cytocinch per cell (right axis, grey) at given time points. ‘n’ denotes the number of buds. p values as labelled (Mann Whitney- U test). (**B**) Individual data points, mean± s.d. and violin plot of cytocinch tether orientation measured with respect to (w.r.t) the longitudinal bud axis at given time points. ‘n’ denotes the number of tethers. p values as labelled (Mann Whitney- U test). (**C**) Bud aspect ratio plotted as a function of cytocinch anisotropy (number*orientation) at different stages. Grey line is a linear fit with R^2^=0.96. (**D**) Schematic of the 2D vertex model with apical, lateral and basal surface tension, enclosing a prescribed bud-ECM volume, with addition of inward forces from circumferential cytocinches (Λ). (**E**) Equilibrium configurations of a bud section, using the 2D vertex model (n=10 cells, grey arrows for basal, lateral and apical surfaces and green dots for cytocinches) at 1 hpb (Λ=0) and 9 hpb (Λ=0.8/cytocinch). (**F**) Predicted evolution of bud radii in the long (blue) and short (orange) axes (experimental: dots, predictions: lines) assuming the experimentally observed increase of bud-ECM volume and cytocinch fraction in time. (**G**) 2D vertex model configuration with and without cytocinch. (**H**) 2D sections of an anterior bud at 6 hpb using Myosin II reporter before and after Cytochalasin D (CytoD 1mM) treatment. Cytocinch is marked by white arrow before treatment. Cytocinch is lost after CytoD treatment. Before (blue) and after (red) circumferential bud diameter is labelled. (**I and J**) Number of cytocinch tethers per bud (I), bud radii and aspect ratios (J) before and after CytoD treatment. ‘n’ denotes the number of buds. p values as labelled (Mann Whitney- U test). (**K**) Illustrations showing the mechanisms underlying budding morphogenesis for SCC formation. Patterned cells in the OV (in orange) synthesize HA. HA drives budding through isotropic extracellular forces (in purple). Anisotropic resistance from cytocinches (in green) mediate anisotropic bud extension.

To experimentally test if cytocinches direct bud extension via mechanical anisotropy, we inhibited their tension using either Cytochalasin D (CytoD), an antagonist of F-actin polymerization, or SMIFH2, an inhibitor of Formin (an Actin polymerization protein). Injecting a mild drug dose (1mM) in the periotic space did not affect OV integrity on short time-scales, but resulted in complete loss of cytocinches and subsequent widening of adjacent membrane angles (Figures 7H, 7I and S6A-S6E). By increasing bud radii in the circumferential axis, loss of cytocinches rendered the aspect ratio of the buds significantly isotropic (Figures 7H, 7J, S6A, S6C, S6D, S6F, Movie S8), showing that cytocinches restrain extension in this axis. Together with theory and experiments, we conclude that biased circumferential orientation of cytocinches constrain the isotropic hyaluronate-pressure into an anisotropic bud extension (Figure 7K, Figure S7, and Movie S8).

## Discussion

Previous studies of tissue morphogenesis typically impute actomyosin contractility as the force producing process. The ECM is deemed as a passive mechanical element that might help shape those forces. Our observations challenge this standard model. In the inner ear, we show that patterned synthesis and swelling of HA-rich ECM creates isotropic pressure to power bud growth. Biased orientation of actomyosin-rich cytocinches restrain growth in the circumferential axis shaping the bud towards a tube. Spatial patterning of hyaluronate-pressure combined with anisotropic tissue material properties from cytocinches may serve as a broadly applicable mechanism for sculpting tissues during development. Remarkably, hyaluronan synthases are locally expressed in a variety of morphogenic epithelia such as the tooth placode, the endocardium and the lens placode (Felszeghy et al., 2001; Haack and Abdelilah-Seyfried, 2016; Kwan, 2014; Tien and Spicer, 2005). Similar to the otic epithelium, HA may be creating pressure to drive remodeling of these tissues. Furthermore, local expression of HA is conserved in the otic epithelial cells of mice (Tien and Spicer, 2005), suggesting a conserved mechanism for SCC morphogenesis across species. The upstream developmental signals that pattern the otic epithelial cells for HA-expression and budding remain to be investigated.

HA is involved in a number of cellular processes including migration, adhesion and differentiation due to binding with cell surface receptors and interactions with other ECM components such as proteoglycans (Chanmee et al., 2015; Cyphert et al., 2015; Toole, 2001). For example, in mouse, covalent modification of HA by the enzyme Tsg-6 is required for mid-gut laterality (Sivakumar et al., 2018). In zebrafish, the bud cells of the otic epithelium express Versican (*vcana* and *vcanb*) (Geng et al., 2013), a proteoglycan that can regulate the structural properties of HA (Wight, 2017). The contribution of HA-mediated signaling and binding with proteoglycans in SCC morphogenesis still remains to be explored. In addition, direct measurements of the physical properties of the HA-rich ECM, i.e. pressure from osmotic swelling and elasticity from interaction with proteoglycans, will be required to disentangle the contribution of each parameter in deforming epithelia. A better understanding of hyaluronate biology will have important implications not just for developmental processes, but also for tissue engineering, regeneration and organoid biology.

Our work shows that actomyosin-rich cytocinches resist hyaluronate-pressure. Importantly, these cinches are tensed and serve as elastic elements that stiffen the tissue selectively in one axis to allow deformation in the orthogonal axis. Most literature on actomyosin attributes its role in morphogenesis to contractility gradients that generate local cell deformations (Salbreux et al., 2012). However, there is an important role of actomyosin networks in stiffening tissues without powering morphogenesis. For instance, actomyosin-dependent tissue stiffening has been observed during axis elongation in frogs and suggested to maintain tissue architecture (Zhou et al., 2009). Developmental biologists tend to equate a high local density of Myosin II with active contractility (Streichan et al., 2018). Our data suggest this assumption should always be questioned. Resistance to forces by anisotropic tissue material properties from cytocinches may serve as a broadly applicable mechanism in tissue morphogenesis.

Interestingly, HA expression is known to induce cellular protrusions in cancer cell lines, thereby enhancing their invasiveness (Kyykallio et al., 2020; Oliferenko et al., 2000; Twarock et al., 2010). In addition, HA expression is essential to epithelial-mesenchymal transition (EMT) during cancer metastasis (Toole et al., 2005). These studies raise an interesting possibility that HA production may be sufficient for cytocinch production. Further investigation is needed to determine how cytocinches relate to other cellular protrusions (Chauhan et al., 2009; Fierro-González et al., 2013; Jacinto et al., 2000; Ramírez-Weber and Kornberg, 1999), and how they are tethered and polarized.

With our work, it is now clear that sources of force production can be cellular (e.g. actomyosin contractility), tissue-scale (e.g. global proliferation), or ECM-derived (e.g. hyaluronate-pressure). Sources of anisotropic stiffness can also be cellular (e.g. cortical tension), tissue-scale (e.g. cytocinches), or ECM-derived (aligned ECM fibres). Our data compels a unifying paradigm for tissue morphogenesis, whereby any or a combination of these sources of force production and anisotropic stiffness can sculpt intricate organs from simple embryonic tissues.

## Acknowledgements

We thank Ian Swinburne, Tony Tsai, Sandy Nandagopal and Toru Kawanishi for support, discussions and reagents. We thank Vanessa Barone, Joseph Nasser and members of the Megason lab for useful comments on the manuscript and general feedback. We are grateful to the Heisenberg lab for transgenic fish. This work was supported by R01DC015478 to S.G.M. A.M. was supported by Human Frontiers Science Program LTF and K99HD098918.

## Author contributions

A.M., T.M. and S.G.M. conceived the project and designed the experiments. A.M. did all the experiments and analysis. E.H. developed the theoretical model. A.M, T.M. and S.G.M. wrote the manuscript. All authors edited the manuscript.

## Competing interests

The authors declare no competing interests.

## Supplemental Figures

**Figure S1.**
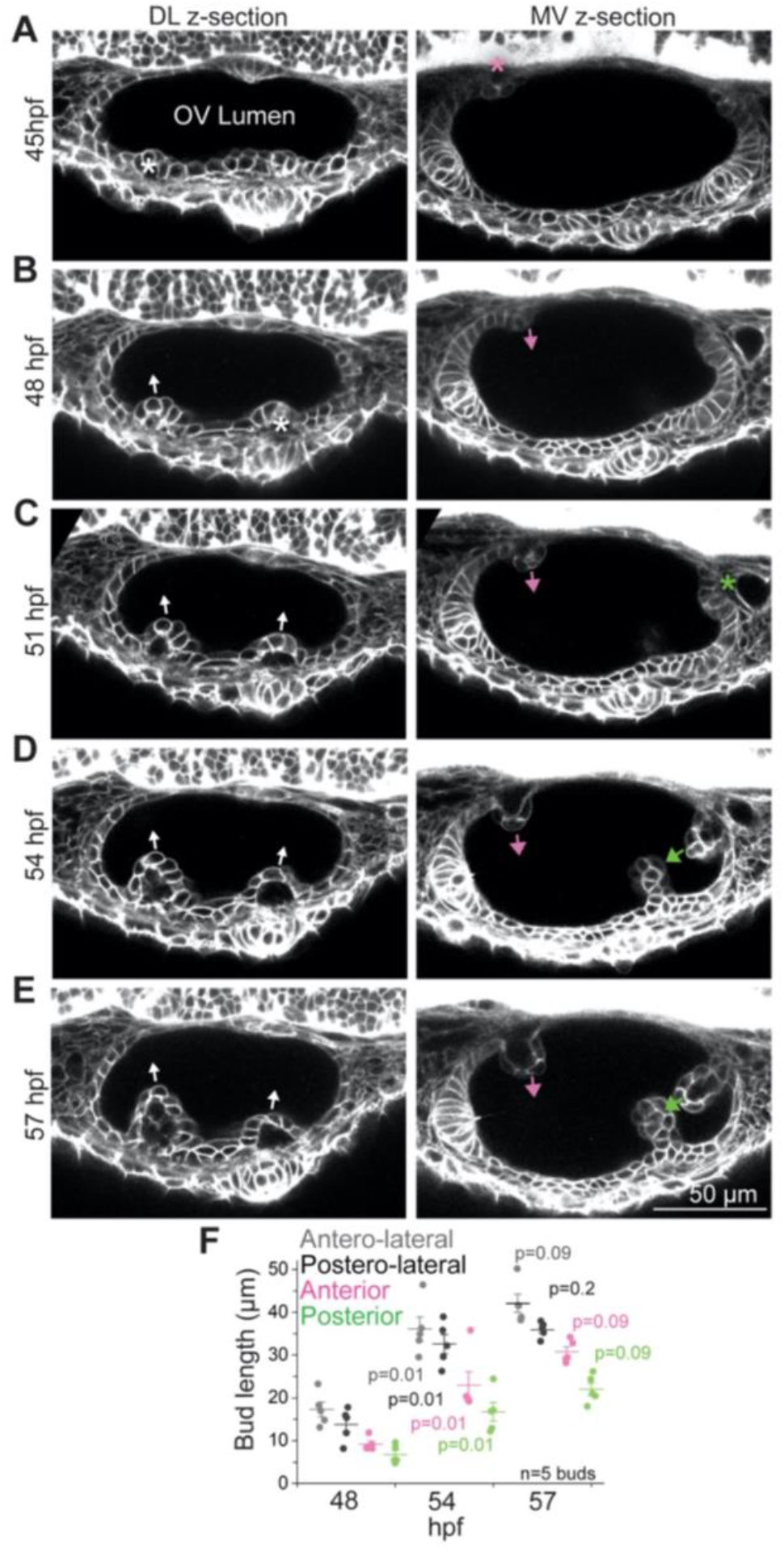
(Related to Figure 1)- *In toto* imaging of the developing zebrafish inner ear reveals multi-scale dynamics during semicircular canal morphogenesis. (**A-E**) Select time points of the OV using *Tg(βActin:membrane-Citrine)* showing 2D sections from two different z-depths. Dorso-lateral (DL) section shows formation of lateral buds (marked by white asterisks) and their extension (white arrows). Medio-ventral (MV) section shows anterior and posterior bud formation (pink and green asterisks respectively) and their extension (pink and green arrows respectively). Scale bar, 50 μm. (**F**) Individual data points and mean±s.d. of bud lengths across time. In the absence of buds, lengths correspond to cell lengths. ‘n’ denotes the number of buds. p values as labelled (Mann Whitney- U test).

**Figure S2.**
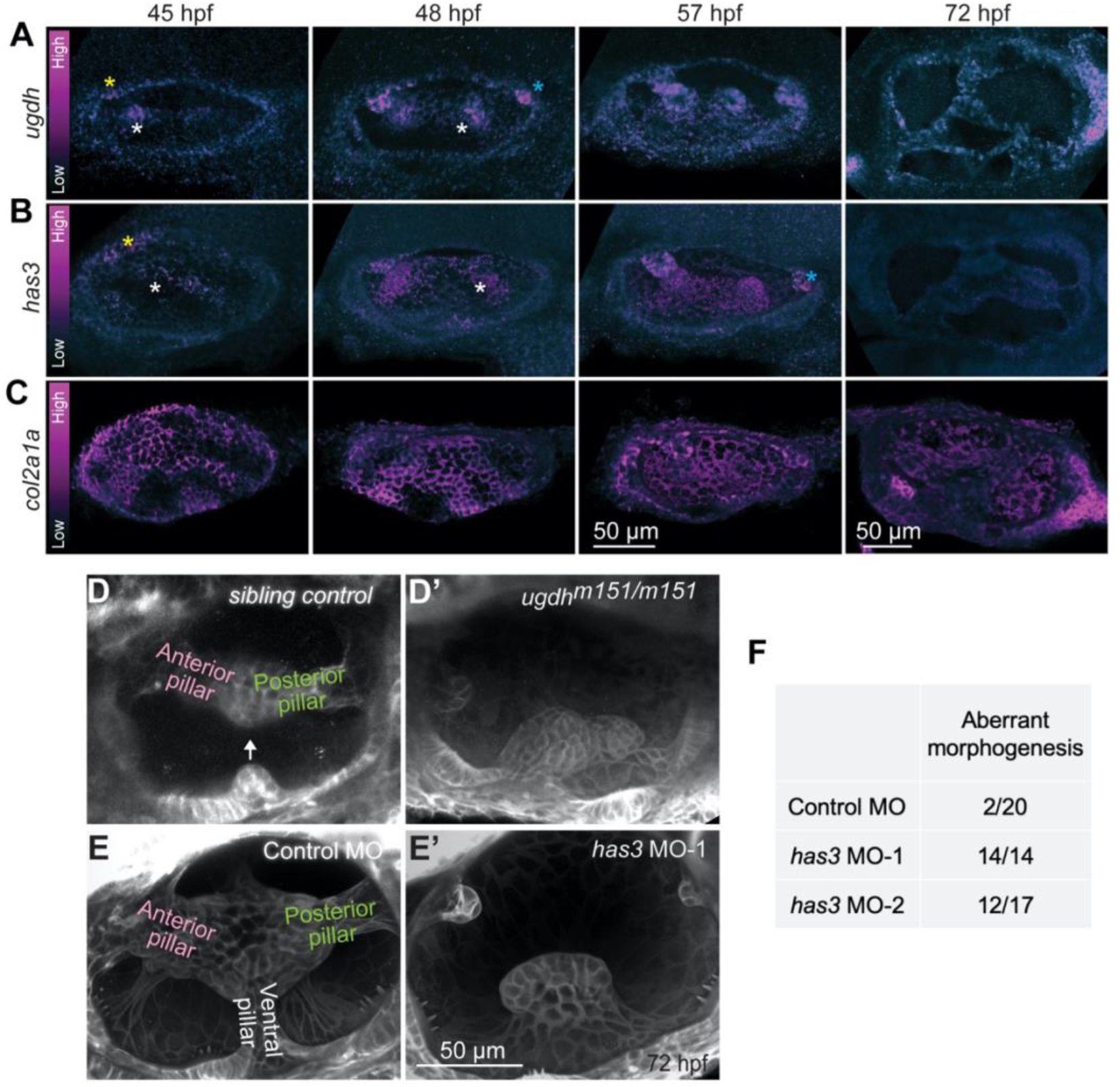
(Related to Figure 3)- Hyaluronan synthesis is required for semicircular canal morphogenesis and patterned through synthesis enzymes *ugdh* and *has3*. (**A-C**), Representative examples of 3D-rendered OVs at select time points, stained with fluorescent *in situ* probes against *ugdh* (A), *has3* (B), and *col2a1a* (C). Heat map represents fluorescent intensity with the same contrast for each probe across time points. Lateral buds are marked with white asterisks. Anterior and posterior buds are marked with yellow and cyan asterisks respectively. Anterior to the left and DL into the plane of view. Scale bar, 50 μm. (**D-E**) Representative examples of 3D-rendered OVs at 72 hpf from sibling control and *ugdh*^*m151/m151*^ mutant larvae labelled with *membrane-NeonGreen* mRNA (D and D’), and *Tg(βActin:membrane-Citrine)* embryos injected with control morpholino (MO) and *has3*-specific MO-1 (E and E’). Anterior and posterior pillars have formed and the ventro-lateral bud is extending (white arrow) in sibling control (D). All three pillars have formed in the control MO (E). Genetically perturbed larvae do not have pillars (D’-E’). Anterior to the left and DL into the plane of view. Scale bar, 50 μm. (**F**) Table showing the number of larvae over the total number measured with aberrant SCC morphogenesis in MO injections at 72 hpf.

**Figure S3.**
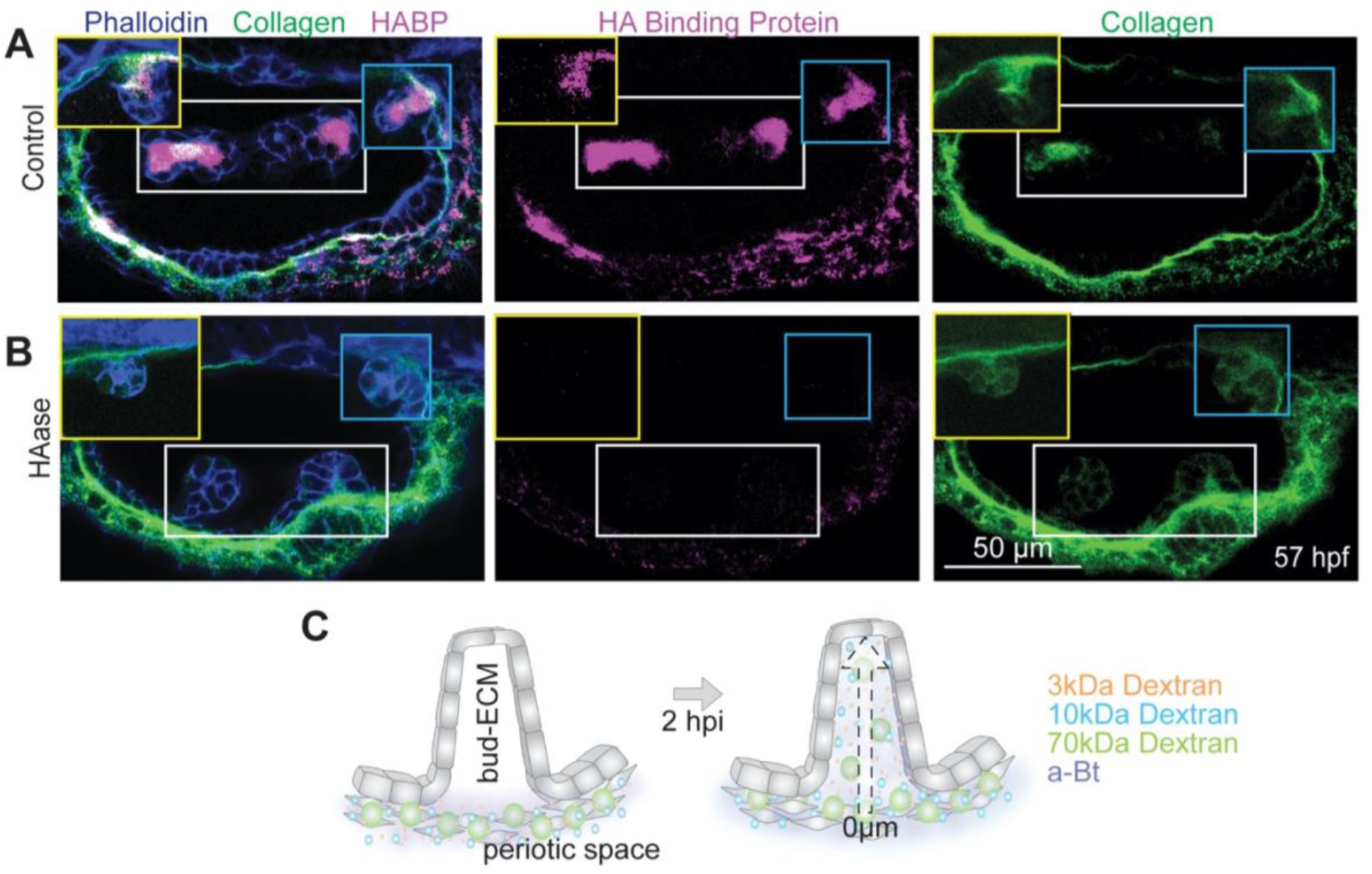
(Related to Figure 4)- HA forms a dense ECM. (**A-B**) 2D sections of OVs showing HA, F-actin and Collagen 2 staining using HA-Binding Protein (HABP), Phalloidin and Anti-Col2a1a respectively, in control and hyaluronidase (HAase) treated larvae at 57 hpf. Lateral, anterior and posterior buds from different z-depths are framed in white, yellow and cyan respectively. Contrast is different at different z depths to capture the dynamic range, and the same across control and treated larvae at the same z-depth. Scale bar, 50 μm. (**C**) Illustration showing different sizes of dextran and alpha-Bungarotoxin (aBt) injected in the periotic space percolating into the bud-ECM.

**Figure S4.**
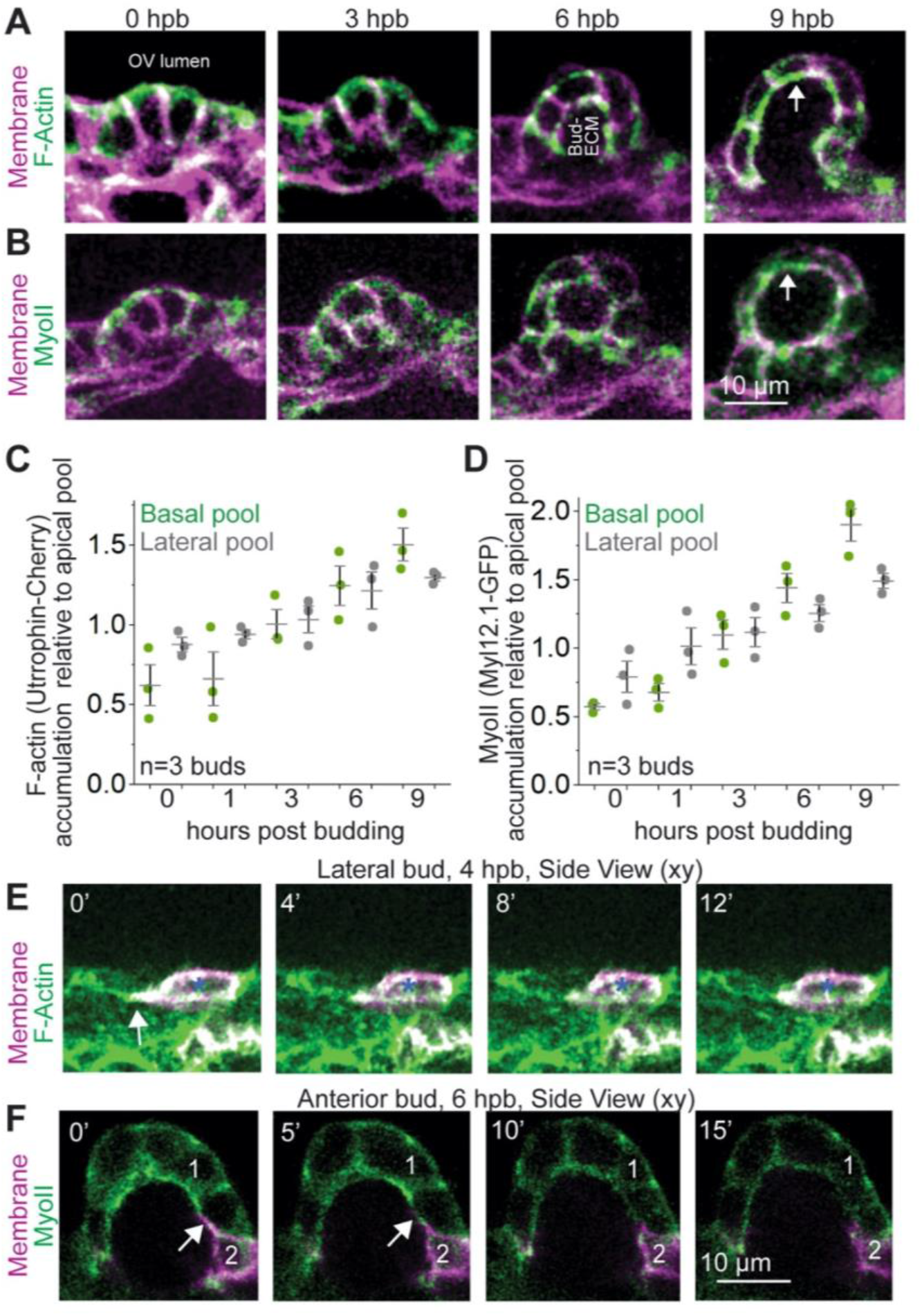
(Related to Figure 6)- Actomyosin networks accumulate on the baso-lateral surface and in tethered membrane protrusions during bud-to-tube transition. (**A and B**) Maximum intensity projections at select time points from posterior buds showing localization of F-actin (using *Tg(βActin:Utrophin-mCherry)*) (A) and Myosin II (using *Tg(βActin:myl12.1-eGFP)*) (B) in green. Membrane (in magenta) is labelled using *Tg(βActin:membrane-Citrine)* in (A) and *Tg(βActin:membrane-mCherry)* in (B). Apical cell surfaces face OV lumen and basal cell surfaces face the bud-ECM. Scale bars, 10 μm. (**C and D**) Individual data points and mean±s.d. of F-actin and Myosin II intensities at the basal and lateral surface normalized to the intensities at the apical surface over time. Each data point is an average from 3-6 cells per bud. ‘n’ denotes the number of buds. (**E**) 2D sections of a lateral bud at select time points showing protrusive activity with mosaic membrane labelling, and the corresponding localization of F-actin (using *Tg(βActin:Utrophin-mCherry)*). Scale bar, 10 μm. (**F**) 2D sections an anterior bud at select time points showing protrusive activity with mosaic membrane labelling, and the corresponding localization of Myosin II (using *Tg(βActin:myl12.1-eGFP)*). Scale bar, 10 μm.

**Figure S5.**
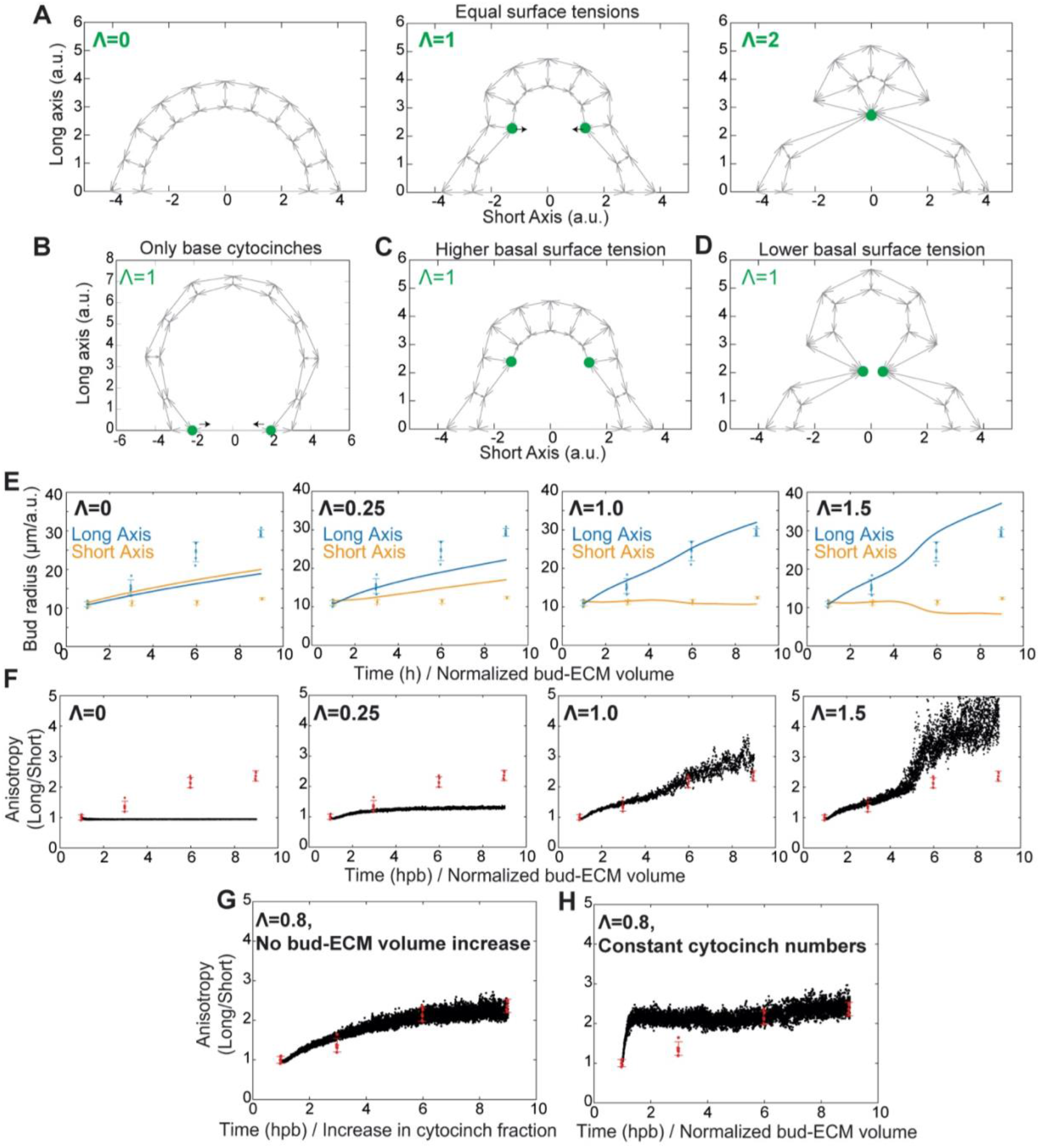
(Related to Figure 7) - A vertex model for bud-to-tube morphogenesis. (**A**) Equilibrium configurations of a section of a bud, modelled via a 2D vertex model (n=10 cells) representing the apical, lateral and basal tensions on the cell surfaces (grey arrows, taken all as equal to unity), as well as inward forces (black arrows) from cytocinches (green dots) with increasing tension (Λ=0, Λ=1 and Λ=2 from left to right panels). (**B**) Equilibrium configurations of a section of a bud with cytocinches only at the base (Λ=1), showing that this is not sufficient to drive anisotropic bud extension, as the monolayer is predicted to be locally spherical (due to isotropic tensions everywhere but at the base). (**C**) Equilibrium configurations of a section of a bud with cytocinches (Λ=1), increased basal tension and decreased apical tension (same parameters as panel a otherwise), showing a better resistance of the epithelium to local deformations. (**D**) Equilibrium configurations of a section of a bud with cytocinches (Λ=1), decreased basal tension and increased apical tension (same parameters as panel a otherwise), showing a worsened resistance of the epithelium to local deformations, and associated bud pinching. (**E and F**) Predicted evolution of the bud radii in the long (blue) and short (orange) axes (experimental: dots, predictions: lines) (E), and their ratios representing bud anisotropy (F) (data in red, model in black, assuming a linear bud-ECM volume increase (3-fold) and temporal increase in fraction of cytocinches (see Methods for details)), for different values of cytocinch tension Λ (left to right panels)). For low cytocinch tension Λ, the bud retains low anisotropy during growth under-estimating experimental anisotropy (left panels), while for high cytocinch tension (Λ=1.5), the model over-estimates bud anisotropy (right panels). Intermediate values of cytocinch tensions provide a good fit for the data (middle panel, Λ=1.0, to be compared to the best-fit value of Λ=0.8 shown in Figures 7E and 7F). Error bars show mean and s.d. (**G and H**) Predicted evolution of bud anisotropies (data in red, model in black), assuming either a constant, non-varying ECM volume and temporal increase in fraction of cytocinches (see Methods for details) for intermediate cytocinch tension Λ=0.8 (G), or a linear bud-ECM volume increase in time (3-fold) and constant fraction of cytocinches (see Methods for details) for intermediate cytocinch tension Λ=0.8 (H). Error bars show mean and s.d.

**Figure S6.**
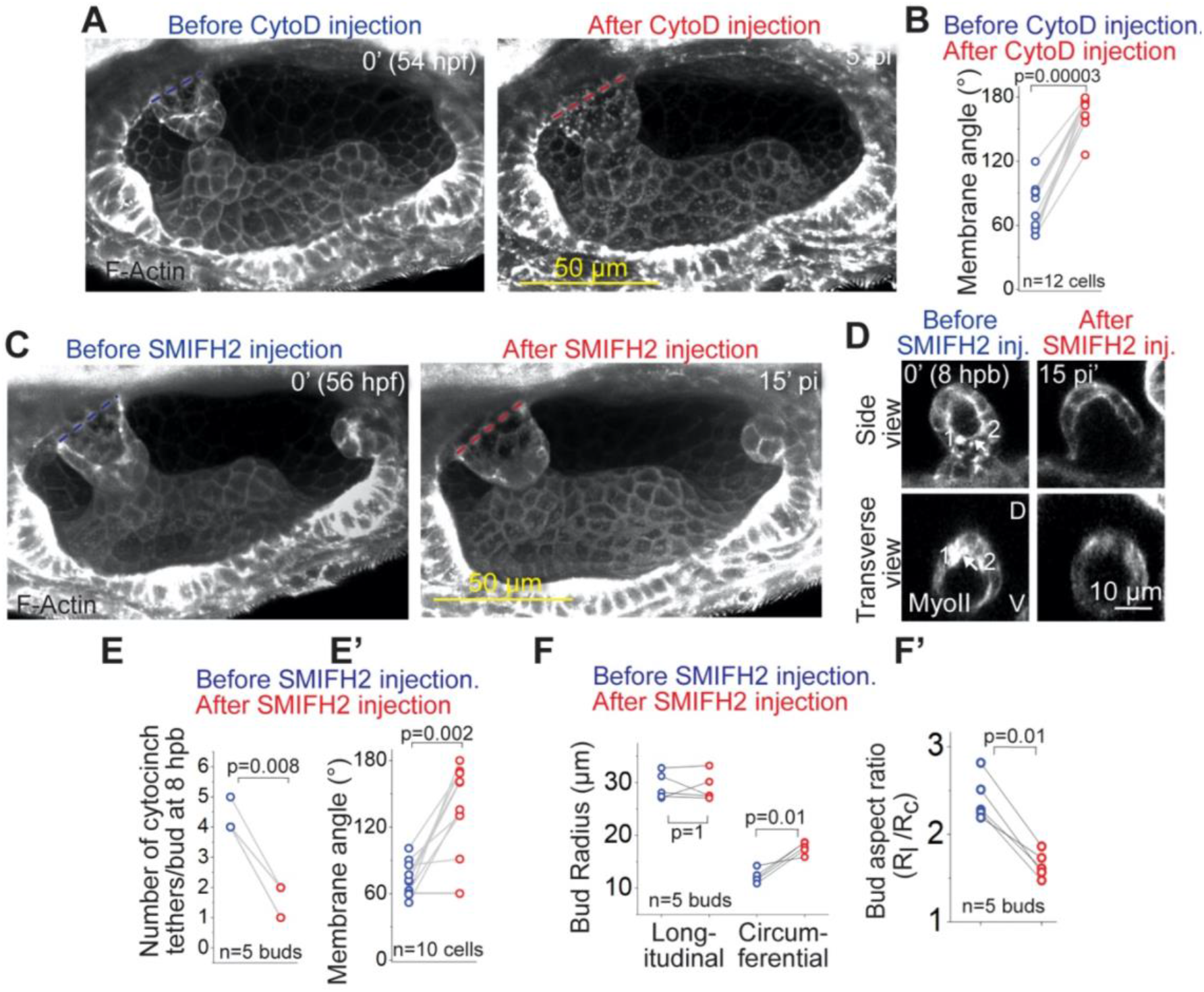
(Related to Figure 7)- Anisotropic stiffness from cytocinches is required for bud-to-tube morphogenesis. (**A and C**) 3D-rendered OVs using *Tg(βActin:Utrophin-mCherry)* at 54 hpf before and after Cytochalasin D (CytoD 1mM) treatment (A), and SMIFH2 (1mM) treatment (C). Anterior to the left and into the plane of view. Scale bar, 50 μm. (**B**) Adjacent membrane angle change before and after CytoD injection. ‘n’ denotes the number of cell membrane angles. p values as labelled (Mann Whitney- U test). (**D**) 2D sections of an anterior bud at 8 hpb using *Tg(βActin:myl12.1-eGFP)* before and after SMIFH2 treatment. Cytocinch is marked by white arrow before treatment. Cytocinch is lost after treatment. Scale bar, 10 μm. (**E and E’**) Total number of cytocinch tethers (E) and adjacent membrane angles (E’) before and after SMIFH2 treatment. ‘n’ denotes the number of buds. p values as labelled (Mann Whitney- U test). (**F and F’**) Bud radii (F) and aspect ratios (F’) before and after SMIFH2 treatment. ‘n’ denotes the number of buds. p values as labelled (Mann Whitney- U test).

**Figure S7.**
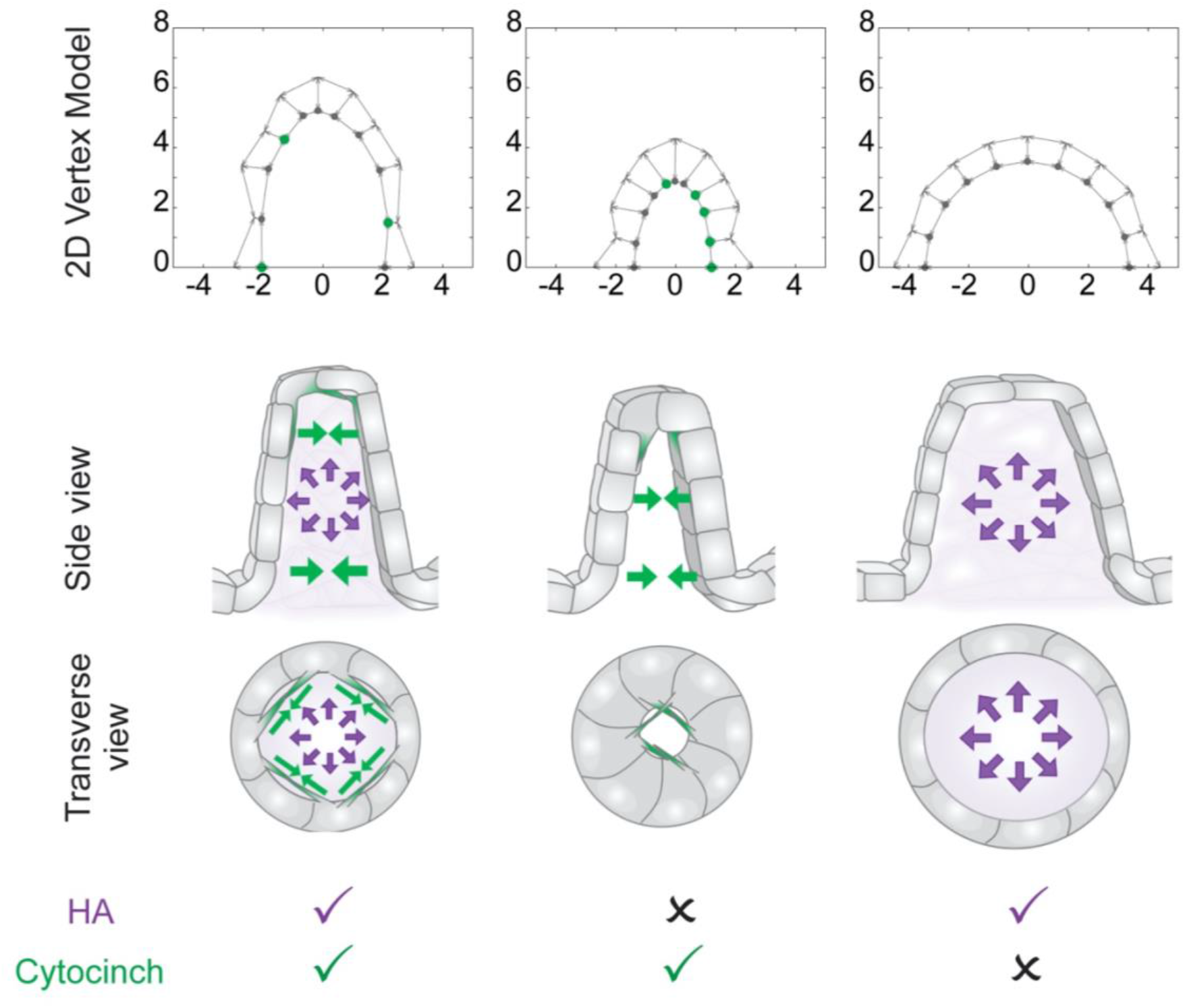
(Related to Figure 7)- Model and experiments show that isotropic hyaluronate pressure shaped by anisotropic stiffness from cytocinches drives semicircular canal morphogenesis. 2D vertex model (top) and illustrations (bottom) showing the effects of HA-mediated isotropic extracellular pressure (purple arrows) and cytocinch-mediated anisotropic tissue resistance (green arrows) on bud and cell morphologies in control and perturbed conditions.

## Experimental Procedures and Resources

### 1) Resource Availability

Information and requests for materials, reagents, resources, data, and code should be addressed to the Lead Contact, Sean Megason.

### 2) Experimental Model and Subject Details

Zebrafish (*Danio rerio)* AB wild-type strain was used. Adult fish were kept at 28.5°C on a 14 hours light/ 10 hours dark cycle. Embryos were collected by crossing female and male adults (3-24 months old). Fish husbandry and fish-related experimental procedures were carried out with approved guidelines from the Animal Welfare Assurance on file with the Office of Laboratory Animal Welfare. The Assurance number on file is #D16-00270.

### 3) Methods Details

#### Zebrafish strain and lines

1. Transgenic- *Tg(βActin:membrane-Citrine): Tg(actb2:mem-citrine-citrine)*^*hm30*^ (Megason lab (Xiong et al., 2013)) used in Figures 1D-1M, S1A-S1E, 2G, 2H, 3E, S2E, 4F-4H, 5A, 5B, 5G, S5A, S5E, 6C and Movies S1 to S6.
2. AB wildtype: Figures 2A and 2B
3. Pigmentation triple mutant- ((*roy*^−/−^; *nacre*^−/−^; *fms*^−/−^) (Parichy et al., 2000; White et al., 2008)) used in Figures S2A-S2C, 3A, 3B, S3A and S3B. The triple mutant lacks all pigmentation rendering the OVs optically clear for fluorescent imaging.
4. *ugdh* mutant: ugdh^*m151/m151*^ is a missense point mutation that disrupts *ugdh* (Driever et al., 1996; Walsh and Stainier, 2001) used in Figures 3D and S2D. Homozygous mutants (25%) were screened from heterozygous parents through characteristic pericardial edema phenotype^22^. Unaffected sibling embryos (75%) could either be homozygous wild-type (50%) or heterozygous for the mutation (25%).
5. Transgenic- *Tg(βActin:Utrophin-mCherry):* Tg*(actb1:mCherry–utrCH)* (Heisenberg lab (Behrndt et al., 2012)) used in Figures 6A (top), 6B (top), S4A, S4E, S6A, S6C and Movies S4 and S5.
6. Transgenic- *Tg(βActin:Myl12.1-eGFP): Tg(actb2:myl12.1-EGFP)*^*e2212*^ (Heisenberg lab (Behrndt et al., 2012)) Figures 6A (bottom), 6B, S4B, S4F, S6D, 7H and Movies S4 and S5.
7. Transgenic- *Tg(βActin:membrane-mCherry): Tg(actb2:mem-mcherry2)*^*hm29*^ (Megason lab (Xiong et al., 2014)) in Figures 2E, 2F, 6B (bottom), 6F (tethered), S4B, and Movie S4).

#### Construct generation and injection of mRNAs, plasmids and morpholinos

The a*ctb2:mem-neongreen-neongreen* construct was made using previously reported membrane localization tags with two copies of moxNeonGreen subcloned downstream of the *actb2* promoter into a pMTB backbone (described here (Collins et al., 2018) in detail).

mRNA was synthesized from a linearized plasmid using the mMessage mMachine T7 transcription kit (Ambion) and injected in single cell stage embryos using a Nanoject system calibrated to 2.3nl of injection volume. *ugdh* mutant embryos were injected with 90pg of *membrane-neongreen* mRNA (Figures 3D and S2D). For mosaic membrane labelling, 90 pg of Tol2 mRNA was injected with either a*ctb2:mem-neongreen-neongreen* or *actb2:mem-mcherry2* (15ng/ul final concentration) into embryos from either *Tg(βActin:Utrophin-mCherry)* or *Tg(βActin:Myl12.1-eGFP)* respectively (Figure 6A, 6B(bottom), 6F (untethered), S4E, S4F and Movie S5).

Two previously characterized morpholino oligonucleotides (MO) (Ouyang et al., 2017) targeting either *has3* start codon (MO-1) or the intron 2-exon 3 splice junction (MO-2) were purchased from Gene Tools along with their Standard Control MO. 7ng of either *has3* MO-1 or *has3* MO-2 or control MO was injected with 3.5ng of p53 MO (to control for phenotypic variability (Gerety and Wilkinson, 2011)) at single cell stage (Figures 3E and S2E).

#### Confocal Imaging

Dechorionated larvae were mounted dorso-laterally (as shown in Figure 1C) using a canyon mount cast in 1% agarose filled with 1X Danieau buffer as previously described in detail (Swinburne et al., 2018). Confocal z-stacks were acquired with an upright Zeiss LSM 710 laser scanning confocal microscope using a C-Apochromat 40X 1.2 NA objective. For single time points, larvae were immobilized by soaking in 1% tricaine. For long-term time lapse imaging, larvae were immobilized with 500 μM α-bungarotoxin protein (aBt from Tocris) (Swinburne et al., 2015) injected into the heart ~30 minutes prior to imaging (at variable stages depending on the experiment). Time-lapse imaging took place in a home-built incubator at 28.5°C. In a typical experiment, ~600 μm X 600 μm X 150 μm volume with a voxel size ~0.2 μm X 0.2 μm X 0.5 μm captured the entire OV within ~3 minutes. The volume and speed of acquisition was adjusted depending on the experiment.

#### Two-photo laser ablation

Ablation of cells in the OV were performed using a Mai-Tai HP 2-photon laser (Spectra-Physics, Santa Clara, CA) as previously described in detail (Mosaliganti et al., 2019). Briefly, the 2-photon laser was tuned to 800 nm at 50-75% power. The pinhole was opened completely and a spot scan at the target cell was performed for 10,000 cycles. To monitor ablation, 3 kDa Texas red-Dextran dye was injected into the heart to label the periotic space. Successful ablation disrupted the otic epithelium causing the dye to enter the OV lumen.

#### Edu Staining, and Hydroxyurea and Aphidicolin drug treatments

45 hpf dechorionated AB wildtype larvae were kept in 500 uM 5-ethynyl-2′-deoxyuridine (EdU), a proliferative marker (Salic and Mitchison, 2008), without (control), or with 20 mM Hydroxyurea (Thompson Coon, 2010) and 300 μM Aphidicolin (DNA replication inhibition (Ikegami et al., 1978)), and a final concentration of 10% Dimethyl sulfoxide (DMSO), in 1X Danieau buffer on a shaker at 4°C for 1 hour. The larvae were transferred back at 28.5°C in fresh 1X Danieau buffer containing 50 uM EdU, without (control) or with 20 mM Hydroxyurea and 300 μM Aphidicolin, and a final concentration of 1% DMSO for 9 hours (45-54 hpf). Larvae were fixed with 4% paraformaldehyde (PFA) in 1X phosphate buffered saline solution (PBS) at room temperature (RT) for 2 hours. Larvae were then permeabilized in pre-chilled acetone at −20°C for 7min, rinsed in 1% Triton in 1X PBS for 5 minutes (thrice) and blocked with 2% Bovine Serum Albumin (BSA) in 1X PBS for 30 minutes. For EdU detection, larvae were transferred to a freshly prepared click-iT® reaction mixture (Thermo Fisher Scientific) for 30mins, at room temperature (RT) in dark. Larvae were washed with 0.1% Tween-20 in 1X PBS (PBT) for 5 mins (5 times) and stained overnight at 4°C with 4′,6-diamidino-2-phenylindole (DAPI) to detect DNA.

#### Multiplex in situ hybridization chain reaction (HCR)

HCR probes, amplifiers, and hybridization, wash and amplification buffers were bought from Molecular Instruments (Choi et al., 2018). Dechorionated larvae at various stages were fixed with 4% PFA in 1X PBS at RT for 2 hours, followed by PBS washing for 5 mins (five times) and permeabilization in pre-chilled acetone at −20°C for 7 min. Larvae were washed with PBS again, and pre-hybridized in hybridization buffer, followed by incubation in hybridization solution containing 1 pg of probes overnight at 37°C. Larvae were then washed in wash buffer followed by washes with 0.1% of Tween-20 in 5X SSC buffer (5X SSCT) for 5 minutes, twice. Larvae were then incubated in amplification buffer for 30 minutes at RT. Meanwhile, hairpin mixtures were prepared by heating 12 pmol of hairpin 1 and 2 per sample to 95°C for 90 seconds, and snap-cooled in the dark. The prepared hairpins were added to the amplification buffer. Larvae were incubated in the hairpin mixture overnight, at RT in dark. Larvae were then washed in 5X SSCT (5 times) and mounted for confocal microscopy (or stored at 4°C). For each larva, two HCR probes were used and visualized with amplifiers conjugated with AlexaFlour™ (AF) 546 and AF647.

#### Hyaluronan Binding Protein, Phalloidin and Collagen 2 staining

Dechorionated larvae (either uninjected or treated as described below) at various stages were fixed with 4% PFA in 1X PBS at RT for 2 hours, followed by PBS washing for 5 mins (thrice), and permeabilization in pre-chilled acetone at −20°C for 7 min. Larvae were then rinsed in 1X PBT for 5 minutes, thrice, and blocked with 5% BSA in 1X PBT for 60 minutes. Larvae were then incubated with primary solution containing Collagen 2 antibody (from GeneTex, polyclonal, rabbit, 1:200 dilution) and biotinylated-Hyaluronan Binding Protein (HABP from EMD Millipore, 1:100 dilution) at 4°C, overnight. Larvae were washed with 1X PBS (Tween-20 was avoided as HABP binds to HA with lower affinity compared to antibody-immunogen binding) for 15 mins, thrice. Larvae were incubated with a cocktail of fluorescent labelled secondary antibody (against rabbit), Streptavidin (against Biotin) and Phalloidin (to label F-Actin) at 4°C, overnight. Larvae were washed with PBS for 15 mins, thrice and mounted for confocal microscopy (or stored at 4°C).

#### Hyaluronidase, Cytochalasin-D and SMIFH2 treatments

Dechorionated larvae (at various stages) were soaked in 1X tricaine and immobilized in a 1% canyon mount filled with 1X Danieau buffer. All injection solutions were made with 1X PBS/0.5% Phenol Red. *Streptomyces* hyaluronidase (HAase from Sigma dissolved in 1X PBS, 300 unit/ml from 3000unit/ml stock) was injected in the periotic space (anterior and posterior to the OV). Control injection solution did not contain HAase. To monitor injections, 3 kDa Texas-red Dextran (from Thermo Fisher dissolved in 1X PBS, 0.5 mg/ml from 20 mg/ml stock) was injected with HAase or control. Cytochalasin D (from Tocris suspended in DMSO, 1 mM from 10 mM stock) (Burke, 1978) or SMIFH2 (from EMD Millipore suspended in DMSO, 1 mM from 10 mM stock) (Rizvi et al., 2009) or DMSO control (1:10 dilution) were injected in the cardiac chamber.

#### Dextran Percolation Assay

Dechorionated larvae (at 50 hpf) were soaked in 1X tricaine and immobilized in 1% canyon mount cast filled with 1X Danieau buffer. All injection solutions were made with 1X PBS/ 0.5% Phenol Red. Neutral Texas-red dextran (either 3 kDa, 10 kDa or 70 kDa from Thermo Fisher dissolved in 1X PBS, 500 μM from 10 mM stock) was co-injected with AF647 conjugated aBt (from Thermo Fisher dissolved in 1X PBS, 500 μM from 1 mM stock) into the periotic space (anterior and posterior to the OV). Larvae were imaged 2 hours post injection.

### 4) Quantification, Modeling and Statistical Analysis

#### Quantification

The software used for image visualization and analysis were- Fiji (Schindelin et al., 2012), ITK-SNAP (Yushkevich et al., 2006) and FluoRender (Wan et al., 2012).

##### 3D visualization

FluoRender open source software was used for 3D renderings. Monochrome heatmap represents z-depth with darker greys into the plane of view in Figures 1D-1K, 2A, 2B, 2F, 3D, 3E, S2D, S2E, S6A, S6C, Movie S1, S3 and S6. Fire heatmap represents fluorescent intensities in Figures S2A-S2C and 4A-D. Image analysis was not performed on 3D renderings.

##### Bud length measurements

Bud lengths were measured manually using the “straight-line” tool in Fiji freeware software. Lines were drawn in the middle sections of the bud and averaged from three different planes for each bud. Where budding had not started or in perturbations where budding was affected, bud lengths correspond to average cell lengths distinguished by their unique taller morphology (visible in Figures S1A-S1C).

##### Volume measurements

Manual segmentations of the OV lumen volume and bud-ECM volume were performed in ITK-SNAP to create meshes. The number of voxels that belong to the mesh were converted to picolitre using the known voxel resolution.

##### EdU positive nuclei

Total number of nuclei and EdU positive nuclei were manually counted in a fixed volume of 3D rendered OVs (z depth 80 μm starting from the dorsal most plane) in Fiji.

##### Aberrant morphogenesis

Number of larvae injected with either *membrane-NeonGreen* mRNA or morpholinos with visible aberrant morphologies by the total number of larvae measured at 72 hpf.

##### Intensity analysis of multiplex in situ HCR

Fluorescent intensity measurements of the two HCR probes per larva were made in Fiji using maximum intensity z-projections covering the entire anterior bud at 57 hpf (25 μm z-depth). Using the “freehand-line” tool (width 20 pixels), region of interest (ROI) was drawn manually as illustrated in Figure 3C, and the mean intensity traces across the line were obtained using the “plot profile” tool. Larvae were carefully staged and mounted, and lines of equal length were drawn and registered at the first cell of the bud to allow comparison between larvae. Intensity traces were normalized to the maximum value, smoothed and averaged.

##### Intensity analysis of Collagen 2 and HABP staining

Fluorescent intensity measurements of Collagen 2 and HABP staining were made in Fiji using maximum intensity z-projections covering both lateral buds at 48 hpf (20 μm z-depth). Using the “freehand-line” tool (width 10 pixels), ROI was drawn manually as illustrated in Figure 4E and the mean intensity traces across the line were procured using the “plot profile” tool. Larvae were carefully staged, and lines of equal length were drawn and registered at the first cell of the antero-lateral bud to allow comparison between larvae. Intensity traces were normalized to the maximum value, smoothed and averaged.

##### Dextran percolation

Percolation of different sizes of dextran was measured in Fiji using maximum intensity z-projections covering all lateral bud-ECM at 52 hpf (10 μm z-depth). ROIs in the bud ECM and the periotic space were drawn manually (as illustrated in Figure S3C). Mean intensity in the red channel (Texas-red Dextran) was normalized to the mean intensity in the farred channel (aBt). Percolation was measured as a ratio of the normalized intensity in the bud-ECM to the normalized intensity in the periotic space.

##### Structural uniformity of the bud-ECM

Structural uniformity of the bud ECM was measured by drawing a line of width 10 pixels the from base to the tip of the bud (as illustrated in Figure S3C). Mean intensity trace across the line was obtained using the “plot profile” tool for various sizes of dextran and HABP staining. Intensity traces were normalized to their mean, averaged and smoothed.

##### Cell diameters and bud radii

Longitudinal cell diameters and longitudinal bud radii were manually measured in Fiji using the “straight-line” tool in the middle XY plane (side view) of either anterior or posterior buds. Line was drawn across the length of the cell for measuring cell diameter, or across the length of the bud for measuring bud radius. Circumferential and radial cell diameters, and circumferential bud radii were measured in the middle XZ plane (transverse view) (as illustrated by the dotted line in Figure 5G) of either anterior or posterior buds. Circumferential and radial cell diameters were measured using the “line” tool by manually drawing a line across the length and the width of the cell respectively. Circumferential bud radii were measured manually by drawing a polygon outlining the apical circumference of the bud. The polygon was fit to an ellipse by Fiji to obtain the radius (since the short and the long axis of the ellipse were comparable).

##### Intensity analysis of F-actin and Myosin II

Fluorescent intensity measurements of F-Actin (*Tg(βActin:Utrophin-mCherry)* and Myosin II (*Tg(βActin:Myl12.1-eGFP))* were made in Fiji using maximum intensity z-projections covering the cells in the middle plane of either anterior or posterior buds (8 μm z-depth). Using the “line” tool (width 5 pixels), ROI was drawn manually inside the cell at the apical, lateral or basal surface to obtain mean intensities.

#### Cytocinch measurements

1. Minimum tether duration was calculated by measuring the time spent by the protrusions in tethered-form before retraction. The start was either the time point when the tether formed (if captured in the time-lapse) or the first frame of the time-lapse.
2. Distance between cells from which tethers originate a time frame before and immediately after tether recoils, were measured manually using the Fiji “line” tool as illustrated in Figure 6E.
3. Membrane angles of cells from which cytocinches originate a time frame before and immediately after tether recoils, were measured manually using the Fiji “angle” tool.
4. To accurately estimate the total number of cytocinch tethers at a given time point, tethers were tracked in time-lapses using their tension signatures (membrane angle and recoil distances as described above).
5. Cytocinch number per cell was calculated by dividing the total number of cytocinches in tethers (i.e. number of tethers in Figure 7A multiplied by 2) by the total number of the cells in the bud (counted manually from 3D renderings) at a given time point.
6. Cytocinch tether angles were measured with respect to the longitudinal axis of the bud using the Fiji “angle” tool. Cytocinches were detected using their tension signatures (membrane angle and recoil distances as described above).
7. Cytocinch anisotropy- Cytocinch anisotropy was calculating by decomposing tether angles into their x (circumferential) and y (longitudinal) components. y component was then subtracted from x, averaged for all cytocinch tethers in a bud at a given time point, multiplied by the total tether numbers at that time point, and plotted against the aspect ratio of the bud at that time point.

#### Theoretical modeling

Here we provide additional information into the modelling approach, parameter fitting, and data-theory comparisons used to understand bud extension during ear morphogenesis.

##### 1) Model derivation

###### a) 2D vertex model without cytocinches

We start by considering the equilibrium of a single cell, under differential tensions on its apical, lateral and basal areas (respectively denoted *γ*_*a*_, *γ*_*l*_ and *γ*_*b*_). The energy of a single cell of height *h*, apical length *r*_*a*_ and basal length *r*_*b*_ then reads (Fouchard et al., 2019; Krajnc et al., 2013; Okuda et al., 2015; Osterfield et al., 2013):

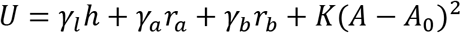

Because the bud that we model are axisymmetric structures, we restrict ourselves here to a 2D vertex description, which will describe well the bud aspect ratio and cellular shapes as assessed by 2D axial sections along the axis of elongation of the bud. Moreover, epithelial cells may be considered incompressible, with constant volume (or here 2D area *A* = *A*_0_, which we implement by taking very large compressibility values *K* → ∞), given the small forces generated by actomyosin structures, compared to osmotic forces (Salbreux et al., 2012). This is valid since cell volume does not change appreciably over the time scales considered (Figure 5C).

We thus consider the ear as consisting of *N* identical such cells (all having the same volume/area and tension parameters *γ*_*l,a,b*_. Finally, given the experimental findings of a key role for ECM-driven pressure for bud growth, we consider that the epithelial monolayer encloses an incompressible fluid of fixed volume *V*_*M*_ (Yang et al., 2020)(which we similarly implement by a quadratic terms around a fixed (but changing in time) area 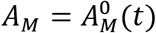 in our 2D description, with high compressibility *K*_*M*_ → ∞), such that the total energy of the monolayer reads

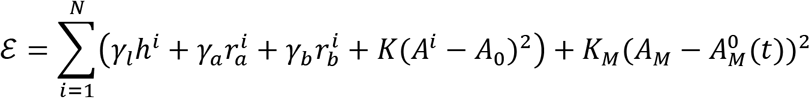

There are therefore two types of control parameters in the system: ECM-controlled temporal changes of volume 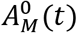 and cellular-level changes in tensions *γ*_*l,a,b*_(largely controlled by actomyosin contractility, which can change the shape of the bud at constant overall volume (Salbreux et al., 2012)).

Finally, this model is complemented with boundary conditions for the border cells *i* = 1 and *i* = *N*, which are the ones remaining in contact with the flat monolayer at position *y* = 0 surrounding the bud. To simulate the rest of the monolayer and ECM attachment, we fix the position of their basal/apical vertex to the plane *y* = 0 while still allowing translations on the plane. Denoting 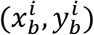 and 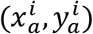 the (*x*, *y*) positions of the basal and apical vertices, this translate into 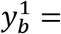 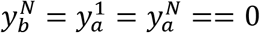. We impose a constant lateral distance for the boundary vertices (as otherwise, the absence of a next cell promotes lateral junction collapse). We also checked alternative, fully clamped boundary conditions (where both *x* and *y* displacements of boundary vertices are forbidden, and found very similar results).

From past theoretical study, we know that such cytocinch-free model will display a buckling transition if there is a mismatch between the preferred area of cells and the volume of the monolayer. For larger hemi-spherical buds (which are indeed observed experimentally when inhibiting cytocinches, see Figures 7H and S6), we can calculate from Eq. S1 that *r*_*a*_ ≈ *r*_*b*_ and the equilibrium configuration of a single cell is 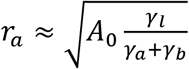, *h* = *A*_0_/*r*_a_. The stress-free state of the bud is one where the ECM volume exactly matches the equilibrium configuration of the monolayer, so that 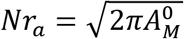, so that for larger 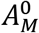, the epithelium will be in a tensile configuration, while being compressed for lower 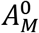, which can trigger monolayer buckling (Figure S5A) (Trushko et al., 2019). Numerical simulations of the equilibrium configurations of the bud for different 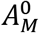 and/or different *γ*_*l*_ confirmed this. Here however, we find no evidence of buckling in the system (see discussion in main text). Instead, ECM volume increase causes a marked flattening of cells, which go from being columnar early on to flattened and squamous like (Figure S4A), as expected in the model for cells with near constant volume. Furthermore, we do not find folding/buckling either upon pharmacological reduction of ECM volume 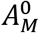 (Figures 5A and 5G), with cells instead going back to cuboidal/columnar, again in agreement with the hypothesis of ECM-driven pressure putting the bud under tension.

Next, we asked whether boundary conditions could be enough to bias the orientation of bud growth, and give rise to elongated structures. We reasoned for instance that a supracellular contractile ring, forming at the base of the buds by cytocinches, could constrain their growth to the axial direction (Turlier et al., 2014). However, in such a model, the local tension within the rest of the monolayer would still be isotropic, which is then predicted to give rise to identical curvatures in both directions (see Figure S5B for example of a numerical simulations with cytocinches only for the vertices at the base), similar to the wetting-dewetting transitions of droplets.

However, given the anisotropy observed in cytocinch tension, we reasoned that this could provide a mechanism to give rise to anisotropic tensions within the monolayer, and therefore proceeded to incorporated them into our model.

###### b) 2D vertex model with cytocinches

We now consider the consequence of cytocinches exclusively in the orthoradial direction (i.e. perpendicular to the 2D section that we model). This induces an additional contribution to surface tension, which tends to shrink locally the radius of the bud (a “purse-string” mechanism), which adds an energy term on basal vertex 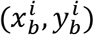 for each cytocinch:

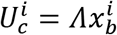

where *Λ* is the cytocinch tension (which tends to minimize the local radius 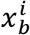 of the bud in the 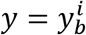 plane), and we note *p*_*c*_ the probability of a given vertex displaying a cytocinch (which increases in time experimentally as well as becoming more directed radially, see Figures 7A and 7B). For the sake of simplicity, we do not include in the model cytocinches in the other, tangential direction, as this simply adds a term which rescales the basal tension *γ*_*b*_.

Before proceeding to run full simulations, we first proceeded to constrain the parameters of the model. From a qualitative perspective, as the bud short-axis is of the order of cell size, and that the tethered cytocinches cause large deflection of vertices (with typical angles *θ*_*m*_ ≈ 90° from Figures 6E-6G), this means from local force balance that cytocinch tension *Λ* should be of comparable magnitude as the surface tensions *γ*_*b*_.

To make this more quantitative, we computed the equilibrium shape of a bud with constant, time-invariant volume and a single cytocinch exerting forces on a single basal vertex (Figures S5A-S5D) - first considering the simplified case of all surface tensions being equal *γ*_*a,l,b*_ = 1. As expected, we found that increasing cytocinch tension increased the local angle of the basal surfaces at the point of force application (Figure S5A), up to a point where cytocinch forces are large enough to pinch off the bud (similar to modelling of cytokinesis). Interestingly, this is something that was observed experimentally, arguing that cytocinch tension is in the high regime.

However, we also found that maintaining *γ*_*a*_ + *γ*_*b*_ constant (and thus the global stretching resistance of the epithelium, see (Yang et al., 2020)), but changing separately the balance of *γ*_*a*_ and *γ*_*b*_ modified the response of the epithelium with respect to local force application. In particular, increased basal tension led to higher bud stability (because this stabilizes the interface on which forces are directly applied, see Figure S5A) whereas increase apical tension led to pinching/collapse of the lateral surfaces (which can occur even for little apical deformation) as shown in Figure S5D. Interestingly, we observe experimentally a relocation of actomyosin from the apical to the basal side in time, as the fraction of cytocinches increases (Figure 7A and S4) - which we conjectured would be an adaptive response giving additional stability to ear bud morphogenesis.

##### 2) Parameter estimation, numerical simulations and predictions

###### a) Parameter estimation and numerical simulations

From these observations and measurements, we then proceeded to constrain further the parameters of the model. After actomyosin relocation (i.e. the relevant anisotropic growth phase with cytocinches), we measured the relative intensities of Myosin on the lateral, basal and apical surfaces (resp. *m*_*l*_, *m*_*b*_ and *m*_*a*_), as a proxy to the tension ratios. We found *m*_*a*_/*m*_*b*_ = 0.4 ± 0.08 and *m*_*a*_/*m*_*l*_ = 0.7 ± 0.3 (see Figure S4).

In the following, we therefore non-dimensionalize all forces by *γ*_*l*_ = 1 and take *γ*_*a*_ = 0.7, *γ*_*b*_ = 1.75. Unless specified, we simulate *N* = 10 cells with varying ECM volume 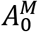 (although we tested different numbers of cells, which did not give rise to significantly different results). We non-dimensionalize all cell length by the preferred cell area 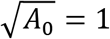 (note that with these values, the equilibrium apical length of a flat cell is 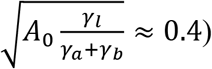, which is close to the experimental value, with an cellular aspect ratio initially around 2: 1 at the early bud stage.

We use a Metropolis algorithm with small temperature to minimize the total energy of the system, performing the simulations multiple times to check that we converge towards the same stead state. Briefly, at each time point, we select *N* times at random a basal vertex at random, move it by a random small change *δr* = 0.05 and calculate the resulting difference in total energy *ΔE* (accepting the move always if *ΔE* < 0 or with a probability *e*^−*ΔE*/*T*^ if *ΔE* > 0 (with *T* = 10^−4^). We then do the same on the adjacent apical vertex. With *K* = 10 and *K*_*M*_ = 0.1, we find that we find all cell areas and ECM area close to their target configuration, while allowing for a non-negligible number of rejected Monte-Carlo steps, so that structures robustly converge to a steady-state for *T* = 10^3^ steps (we checked that doubling the number of steps did not change the results). For numerical stability, we prohibited moves that shrank junctions to values smaller than a small cut-off (*l*_*m*_*in* = 0.3).

Moreover, in simulations where we vary the ECM volume 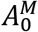, we do so in a slow manner, so that the monolayer is near equilibrium throughout the simulation (*T* = 5.10^4^ steps, although we again checked that doubling the step number did not change the results). Such quasi-static approximation is justified by the fact that ECM growth occurs very slowly at time scale of hours, which are much larger than the timescales of actomyosin-driven shape changes (minutes to tens of minutes). We thus define 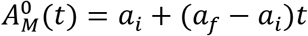 (where we normalized time to be 1 at the end of the simulation, so that *a*_*i*_ and *a*_*f*_ are respectively the initial and final volumes/areas (measured experimentally to be *a*_*f*_ ≈ 3*a*_*i*_ between 1hpb and 9hpb, as shown on Figure 5K).

Furthermore, guided by the increase in frequency of cytocinches in time, to a maximal value of around 0.5 cytocinch per cell (Figure 7A), we define the probability of a cytocinch on a given basal vertex as 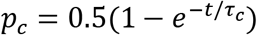 (zero at the beginning of the simulation and 0.5 at the end, with a build-up time of *τ*_*c*_ ≈ 1.5*h* based on Figure 7A, although the results were weakly sensitive to this value). We also tested the effect of a constant, non-time-varying fraction of cytocinches (with *Λ* = 0.8), and found, as expected that although the long-term effect was unchanged, this would predict smaller, early bud to be already highly anisotropic (Figure S5G and S5H), in contrast to the experimental data. This validates our hypothesis that cytocinch (and more specifically the increase of oriented cytocinches as a function of time) is crucial for the growing anisotropy of the buds. On the other hand, we also tested the converse scenario of increasing probabilities of cytocinches (as in WT simulations), but with a constant ECM volume. This led to anisotropies very similar to WT with volume increase (despite long and short axis being much shorter), also validating our hypothesis of ECM-bud volume growth being mainly responsible to overall bud inflation, but not anisotropic growth (Figure S7)

One particular assumption here is that the tension in a given tethered cytocinch is constant in time (and thus that the increased anisotropy of the structure is driven by the increase in cytocinch number), an assumption motivated by the angle of basal deformation caused by cytocinches (Figure 6E-G), which we found to be comparable in early vs. late buds. With this, as well as the various parameters constrained as described above, the main key parameter controlling the bud anisotropy that remains to be fitted is thus the tension *Λ*.

###### b) Comparison with data

For this, we computed the short *a* and long axis *b* of the bud in time during our simulation for different values of *Λ*, as this provides a direct comparison to Figure 5J and 5K. As expected, for *Λ* = 0, the structure was growing spherically, with the ratio of short and long axis converging towards 1 *a/b* → 1 for large volumes, while the anisotropy increased for larger values of *Λ*. Interestingly, we found that for intermediate values of *Λ* ≈ 0.8, the short-axis remained roughly constant in width, while the long-axis increased (3-fold to match the 3-fold increase in volume, see Figure S5E and Movie S8), very close to the experimental observations as shown on Figure 7F. We also plot the anisotropy of the bud in time (calculated as the long axis length divided by the short axis at the base) for different values of *Λ*, overlaying the points from 10 different runs of a simulation with identical parameters, to show the stochastic variability (due to the fact that cytocinch formation is a random event in the model) and robustness.

This behavior at intermediate *Λ* = 0.8 (Figures 4E-4G) contrasts with lower values of *Λ*, for which the short-axis markedly increase in size (Figure S5E) and larger values of *Λ*, characterized in particular by mechanical instabilities which the bud is locally pinched off (and in particular at the base as we have not imposed boundary stresses in the *x* direction). This argues that the data can be well-fitted (Figure 7F) by these intermediary values of constant *Λ* (also consistent with the basal deformation angles observed in the data, as discussed above), and increasing number of cytocinches as a hydrogels function of time. The model also showed near-uniform transition from a columnar epithelium to a squamous one upon bud-ECM volume growth, an effect which stems from the approximation of constant cell volume together with uniform cytocinch distribution in space, and which was reversed upon volume reduction as expected (Figures 5G-5I).

We then proceeded to test this further in the data, by directly comparing cytocinch number and anisotropy ratio *a/b*, with the prediction that these should strongly correlate in a quasi-linear manner (see prediction for different *Λ* on Figures S5E and S5F). Importantly, this was in close agreement with the data (Figure 7C). We also performed several pharmacological experiments to test the model. Disrupting cytocinches via CytoD decreases the aspect ratio *a/b* back to a value close to 1, as a expected for *Λ* ≈ 0 in the model (Figures 7H-7J and Movie S8 for a simulation where we turned off cytocinches to *Λ* = 0 at 8h). Similar behavior was obtained by disrupting cytocinches with SMIFH2 (Figures S6 and S7). Conversely, reducing volume as constant cytocinch strength/numbers is predicted to retain high anisotropy while shrinking (Figures 5G, S5G, S7 and Movie S8 for a simulation where we go back to the initial ECM area *a*_*i*_ at 8h) as experimentally observed upon hyaluronidase treatment which causes a collapse of the ECM (Figure 5G).

#### Statistics

Statistical methods were not used to predetermine sample size. Sample size was chosen such that statistical variability between experiments was captured and did not change significantly when more data points were added. No data was excluded. No experiments were randomized and the investigator was not blinded to any data set. Each experiment was replicated at-least thrice. Statistics were done on all replicates pooled together.

Average values are calculated from ‘n’, where ‘n’ represent number of OVs, buds, bud-ECM, cells, cytocinch tethers, or cytocinches (as labelled in each figure). Plots show individual data points, mean values and error bars (standard deviation or standard error of mean, as labeled). Non-parametric Mann–Whitney *U*-test was used to determine *p* values wherever mentioned. All statistical tests and plots were made in OriginPro.

### 5) Key Resource Table

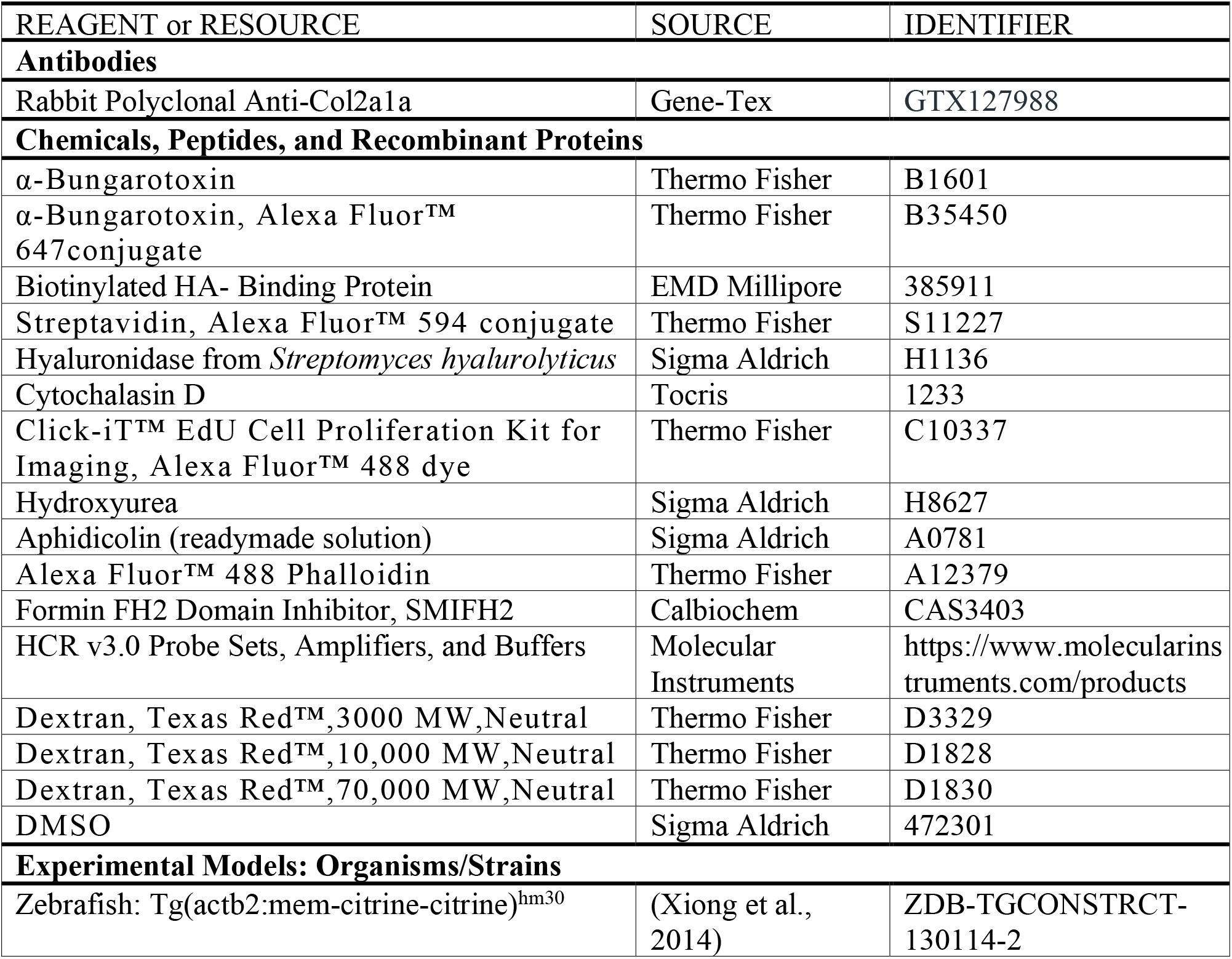
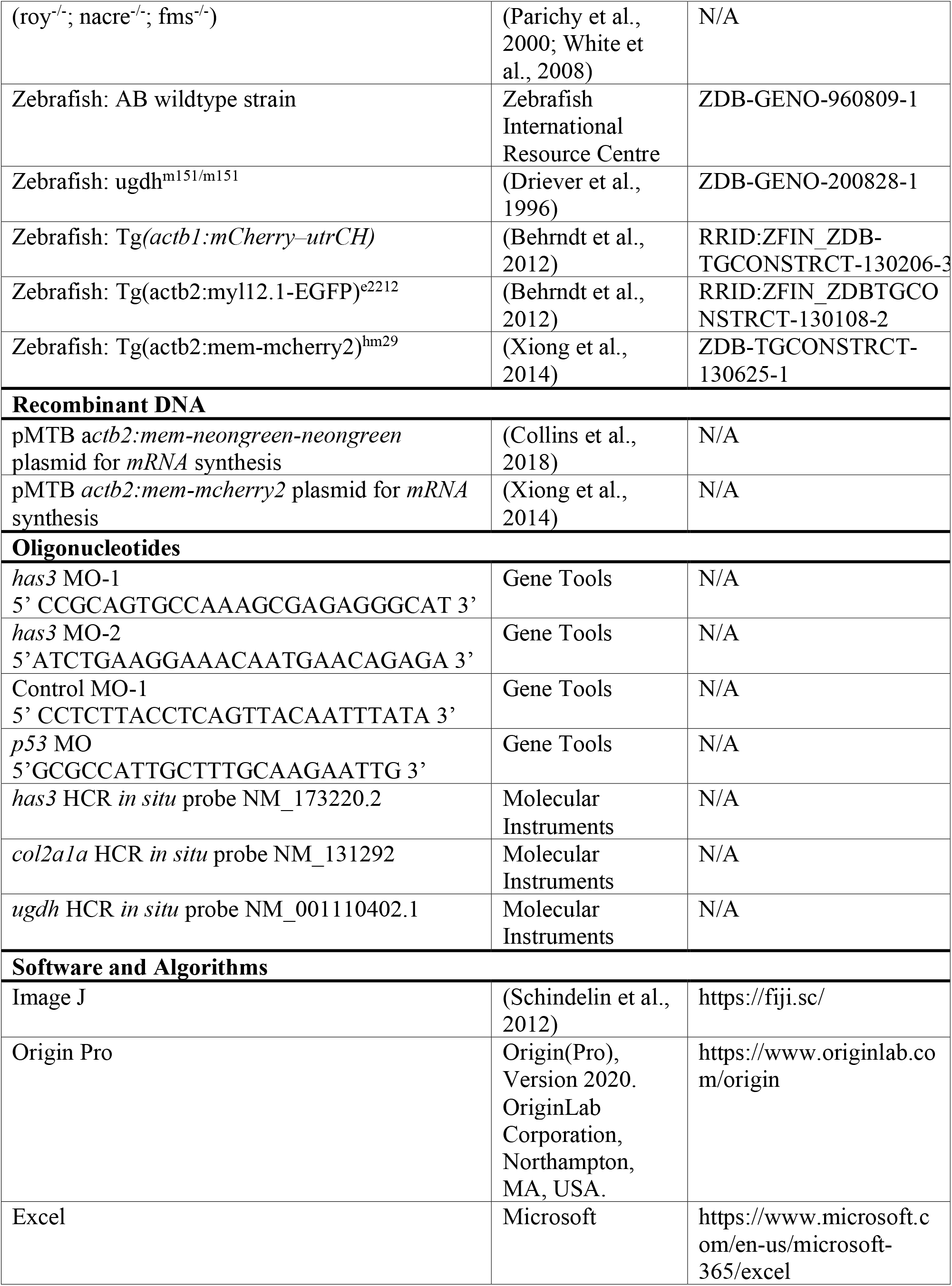
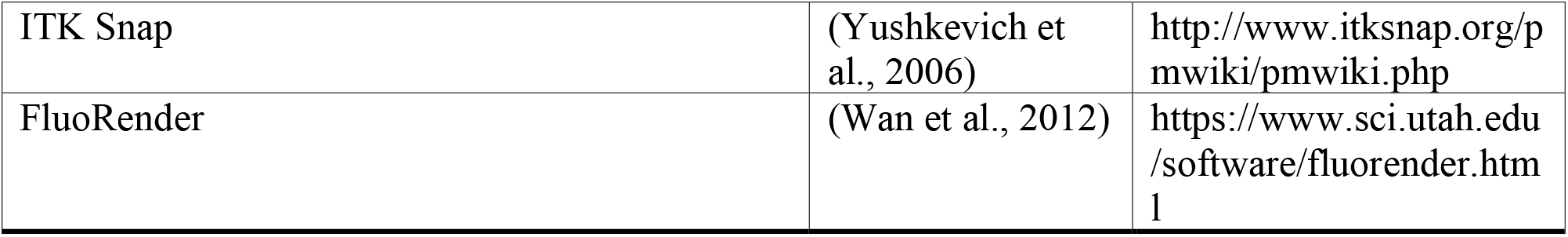

## Supplemental video captions

**Video S1 (Related to Figure 1): Semicircular canal morphogenesis**

3D rendered time-lapse images of the otic vesicle (OV) showing semicircular canal morphogenesis involving bud formation (black asterisks), extension and fusion using two *Tg(βActin:membrane-Citrine)* larvae (change at 59 hpf). OV lumen is coloured (blue) in the first and last frame. Bud fusion demarcates the hubs for the anterior, posterior and lateral semicircular canals (marked by pink, green and white circular arrows respectively). Anterior to the left and dorso-lateral into the plane of view. Scale bar, 50 μm.

**Video S2 (Related to Figure 4): Bud-ECM volume increases during bud extension**

Time-lapse images showing the lateral buds in *Tg(βActin:membrane-Citrine)* larva (green) injected with Texas-red 3 kDa Dextran (magenta) in the periotic space. Scale bar, 20 μm.

**Video S3 (Related to Figure 5): Anisotropic bud and cell morphologies during bud extension**

Side and transverse views of 3D rendered time-lapse images of an anterior bud using (*Tg(βActin:membrane-mCherry*) larva. Longitudinal (blue), circumferential (orange) and radial (white) axes are labelled at 5 hpb. Scale bar, 10 μm.

**Video S4 (Related to Figure 6): Baso-lateral actomyosin localization during bud extension**

Side and transverse views (maximum intensity projections) of time-lapse images showing localization of F-actin (using *Tg(βActin:Utrophin-mCherry)*) and Myosin II (using *Tg(βActin:myl12.1-eGFP)*) in green. Membrane (in magenta) is labelled using *Tg(βActin:membrane-Citrine)* with F-Actin, and *Tg(βActin:membrane-mCherry)* with Myosin II. Scale bar, 10μm.

**Video S5 (Related to Figure 6): Mosaic membrane labelling shows actomyosin-rich membrane protrusions**

2D sections of lateral buds showing protrusive activity (white arrows) with mosaic membrane labelling (magenta), and the corresponding localization of F-actin (using *Tg(βActin:Utrophin-mCherry)*) and Myosin II (using *Tg(βActin:myl12.1-eGFP)*) in green at 4 hpb. Scale bar, 10 μm.

**Video S6 (Related to Figure 6): Cytocinches: Cellular protrusions that form tension-rich tethers**

Side and transverse views of 3D rendered time-lapse images of an anterior bud using (*Tg(βActin:membrane-Citrine)* larva. Tether formation is marked by white arrows. Adjacent membrane angle changes and cell distance is marked in blue and red before and after tether recoil respectively. Scale bar, 10 μm.

**Video S7 (Related to Figure 7): Cytocinches have a polarised distribution biased in circumferential axis**

Side and transverse views (2D sections) of an anterior bud at different depths (0 μm is the most dorsal-lateral plane for side view, and the tip of the bud for transverse view) with cytocinch tethers tracked in white arrows using the Myosin II reporter (*Tg(βActin:myl12.1-eGFP)*). Scale bar, 10 μm

**Video S8 (Related to Figure 7): A 2D vertex model recapitulates control, Hyaluronidase and CytoD treated bud morphologies**

Equilibrium configurations of a bud section over ‘time’ (hours post budding), modelled via the 2D vertex model representing apical, lateral and basal tensions on the cell surfaces (grey arrows), and inward forces from cytocinches (green dots). ECM-bud volume and cytocinch frequency increase in time (as experimentally measured). X and Y axes are short and long respectively. Left panel is control, middle panel represents Hyaluronidase treatment (through reduction in bud-ECM volume), and right panel represents Cytochalasin D treatment (through removal of cytocinches), each at 7 hpb.

## Notes

### Competing Interest Statement

The authors have declared no competing interest.

